# Alkane-oxidizing consortia produce substantial amounts of disaccharides

**DOI:** 10.64898/2026.05.21.726584

**Authors:** Stian Torset, Lennart Stock, Marcus Elvert, Manuel Liebeke, Gunter Wegener

## Abstract

Consortia of archaea and partner bacteria couple the anaerobic oxidation of alkanes to sulfate reduction. While catabolic pathways in anaerobic alkane-oxidizing archaea (ANKA) are increasingly understood, their anabolic capacities remain poorly characterized. Here, we examined nine enrichment cultures dominated by ANKA and their partner bacteria for small-molecular compounds using solvent extraction and gas chromatographic analysis of derivatized extracts. All hydrocarbon-degrading cultures contained substantial amounts of disaccharides in their metabolite pools. Cold-adapted methane-oxidizing cultures dominated by ANME-2c and Seep-SRB2 contained up to 1.5 mg of trehalose per mg soluble protein. Trehalose was also abundant in ethane-oxidizing cultures of Candidatus Ethanoperedens and its distinct partner SRBs, accounting for up to 75 % of the extracted metabolites. In contrast, thermophilic ANKA cultures dominated by ANME-1 or *Ca*. Syntropharchaeum and *Ca*. Desulfofervidus contained an abundant as-yet-unidentified glucose-containing disaccharide. Metagenomic analysis revealed widespread trehalose metabolism genes among partner Desulfobacterota and in ANME-2c and *Ca*. Ethanoperedens, but a lower potential in ANME-1 and Syntropharchaeum, consistent with metabolite profiles. If exogenous trehalose was added to the Ethane50 culture, we observed rapid metabolization by heterotrophic microorganisms, but poor assimilation by the *Ca*. Ethanoperedens/ *Ca*. Desulfofervidus core community, indicating that ANKA/SRB consortia do not consume externally supplied trehalose. Instead, *Ca*. Ethanoperedens/ *Ca*. Desulfofervidus, as well as other ANKA/SRB consortia, may use the disaccharides as energy-storage molecules, osmolytes, or components of the extracellular matrix. Notably, the disaccharides produced by the consortia also sustain ancillary heterotrophs, thereby linking alkane oxidation to broader sedimentary carbon cycling.

## 2 Introduction

In deep marine sediments, methanogenic archaea produce substantial amounts of methane (Duan et al., 2023; Reeburgh, 2007). In addition, the thermogenic decay of biomass and abiogenic reactions between hydrogen and carbon dioxide result in the formation of methane and short-chain alkanes (Sherwood Lollar et al., 2002; Tissot and Welte, 1984). These compounds are the energy substrates for anaerobic alkane-oxidizing archaea (ANKA), which thrive alongside their sulfate-reducing partner bacteria at the sulfate-methane interface (Boetius et al., 2000; Orphan et al., 2001). Among them, the anaerobic methane-oxidizing archaea (ANME), a polyphyletic group, play a critical role in mitigating methane efflux through anaerobic oxidation of methane (AOM) by consuming up to 90% of the methane in marine sediments (about 380 Tg methane per year; Kurth et al., 2019; Martinez-Cruz et al., 2018; Reeburgh, 2007). ANKA also include archaea that oxidize short-to long-chain alkanes, such as the ethane oxidizers of the Candidatus Ethanoperedens (Chen et al., 2019; Hahn et al., 2020), and the propane and butane oxidizers of the *Ca*. Syntropharchaeum (Laso-Pérez et al., 2016), with relatively high abundances at select seeps and hydrothermal vents (Wegener et al., 2022).

All ANKA activate their alkane substrate with variants of the methyl-coenzyme M reductase; most are capable of complete substrate oxidation (Wegener et al. 2022). ANMEs achieve this by a complete reversal of methanogenesis, with only minor modifications for the oxidative directionality of the pathway (Hallam et al., 2004; Thauer and Shima, 2008). The enzymatic chain of non-methane alkane oxidation in archaea is not fully resolved; however, it has been established that due to their lack of a complete respiratory pathway, most ANKA require external electron sinks (Chadwick et al., 2022; Haroon et al., 2013; Knittel and Boetius, 2009; McGlynn et al., 2015). In marine sediments, ANKA form syntrophic consortia with sulfate-reducing bacteria from the phylum Desulfobacterota (Boetius et al., 2000; Murali et al., 2023; Schreiber et al., 2010; Wegener et al., 2015). The transfer of reducing equivalents is most likely mediated by direct interspecies electron transfer (DIET) (McGlynn et al. 2015, Wegener et al. 2015). As exceptions, the freshwater ANME-2d (*Ca*. Methanoperedens) couples reverse methanogenesis to nitrate and metal oxide reduction(Haroon et al., 2013; McIlroy et al., 2023), and the Methanoliparales couples alkane oxidation to methane formation (Zhou et al., 2021). Thus, these archaea do not require partner bacteria for the anaerobic oxidation of alkanes.

Research in the field of anaerobic microbial alkane oxidation focused on the catabolic capabilities of the organisms involved, i.e., how they extract energy from alkanes and how they transfer energy as reducing equivalents to their sulfate-reducing partners (Berger et al., 2021; Chadwick et al., 2022; Hahn et al., 2020; Laso-Pérez et al., 2016; McGlynn et al., 2018, 2015; Timmers et al., 2017; Zehnle et al., 2023). Here, we studied alkane-oxidizing communities’ capacity to produce and metabolize saccharides, focusing on their anabolic pathways and metabolic versatility. We combined metagenomic analysis with metabolomics and found strong accumulation of disaccharides and genomic evidence of disaccharide metabolism in all ANKA/SRB cultures. We also demonstrate, using stable isotope probing, that disaccharides potentially generated by ANKA sustain heterotrophs in the ancillary community.

## 3 Results

### 3.1 Enrichments of ANKA

Here, we analyzed sediment-free, anaerobic alkane-oxidizing cultures derived from distinct marine gas-rich sediments enriched in specific clades of alkane-oxidizing archaea and their partner SRB (Table 1). The low-temperature AOM culture HR12 was initiated from Hydrate Ridge sediment (Cascadia Margin, 780 m water depth) at a cultivation temperature of 12 °C (Holler et al., 2009). The Elba20 culture originated from methane-percolated coastal sands near the island of Elba and was cultured at 20 °C. The Isis20 enrichment was established at 20 °C from sediments collected from the Isis Mud Volcano at 1250 m depth in the Eastern Mediterranean (RV L’Atalante, 2003). It contained a more diverse AOM community, including ANME-2c, ANME-2a, and ANME-2b, associated with Seep-SRB1a (Schreiber et al. 2010). The thermophilic AOM culture GB50 was produced from hydrothermally heated sediments of the Guaymas Basin (AT15-45 expedition, 2009) at 50 °C, and is dominated by ANME-1a and *Ca*. Desulfofervidus auxilii (Krukenberg et al., 2018). Additionally, we investigated a low-temperature ethane-oxidizing enrichment (Ethane20) and a high-temperature ethane-oxidizing enrichment (Ethane50) produced from the Guaymas Basin (AT37-06 expedition 2016; (Hahn et al. 2020) and propane and butane-oxidizing enrichments (Propane50 and Butane50), likewise produced from Guaymas Basin sediments (AT15-45 expedition, 2009) (Laso-Pérez et al. 2016).

**Table 1:** Enrichments and maintenance conditions of alkane-oxidizing consortia obtained from different marine sedimentary environments during various research expeditions.

| Enrichment | ANKA/SRB | Substrate | Temp (°C) | Expedition | Year | Location | Coord. | Depth (m) | Publication |
| --- | --- | --- | --- | --- | --- | --- | --- | --- | --- |
| **HR12** | ANME-2c/Seep SRB 2 | Methane | 12 | SO148-1 | 2000 | Cascadia Margin, NE Pacific | 44.57N, 125.145 W | 780 | Holler *et al.*, 2009 |
| **Elba 20** | ANME-2c/Seep SRB 2 | Methane | 20 | Hydra Diving team | 2010 | Elba | 42.7438 N, 10.1182 33E | 12 | Krukenberg *et al.*, 2018 |
| **ISIS20** | ANME-2c/2a Seep-SRB1a | Methane | 20 | Nautinil Expedition RV L' Atalante | 2003 | ISIS Mud Volcano | 32.350N 31.383E | 1250 | Schreiber *et al.*, 2010 |
| **GB37** | ANME-1a/Seep SRB2 | Methane | 37 | AT15-45 | 2009 | Guaymas Basin | 27.01 N, 111.409 133W | 1999 | Krukenberg *et al.*, 2018 |
| **GB50** | ANME-1a/ *Ca. D. auxilii* | Methane | 50 | AT15-45 | 2009 | Guaymas Basin | 27.01 N, 111.409 133W | 1999 | Krukenberg *et al.*, 2018 |
| **Ethane20** | Ethanoperedens unspc./ DSS partner | Ethane | 20 | AT37-06 | 2016 | Guaymas Basin | 27.008N 111.409 W |  | Unpublished |
| **Ethane50** | *Ca. Ethanoperedens thermophilum* | Ethane | 50 | AT37-06 | 2016 | Guaymas Basin | 27.008N 111.409 W |  | Hahn *et al.* 2020 |
| **Butane50** | *Ca. Syntropharchaeum butanivorans* / *Ca. D. auxilii* | Butane | 50 | AT15-45 | 2009 | Guaymas Basin | 27.01 N, 111.409 133W | 1999 | Lazo-Perez *et al.*, 2018 |
| **Propane50** | *Ca. Syntrophoarchaeum caldarius* / *Ca. D. auxilii* | Propane | 50 | AT15-45 | 2009 | Guaymas Basin | 27.01 N, 111.409 133W | 1999 | Lazo-Perez *et al.*, 2018 |

### 3.2 Disaccharides are abundant in alkane-oxidizing enrichment cultures

We extracted small polar molecules from the above-described alkane-oxidizing cultures and derivatized the extracts for GC-MS-based metabolite analysis (Sogin et al., 2019) (Figure 1 A-E). In all cultures except the ANME-2a/2b/2c-dominated ISIS20, disaccharides were the most abundant metabolites, often accounting for >50% of the total signal (Figure 1A, Figure 1E, Supplementary Tables 1-2). In addition, the metabolite extracts contained derivatized fatty acids and amino acids, such as glutamic and aspartic acids (Supplementary Table 1, Supplementary Table 2).

**Figure 1.**
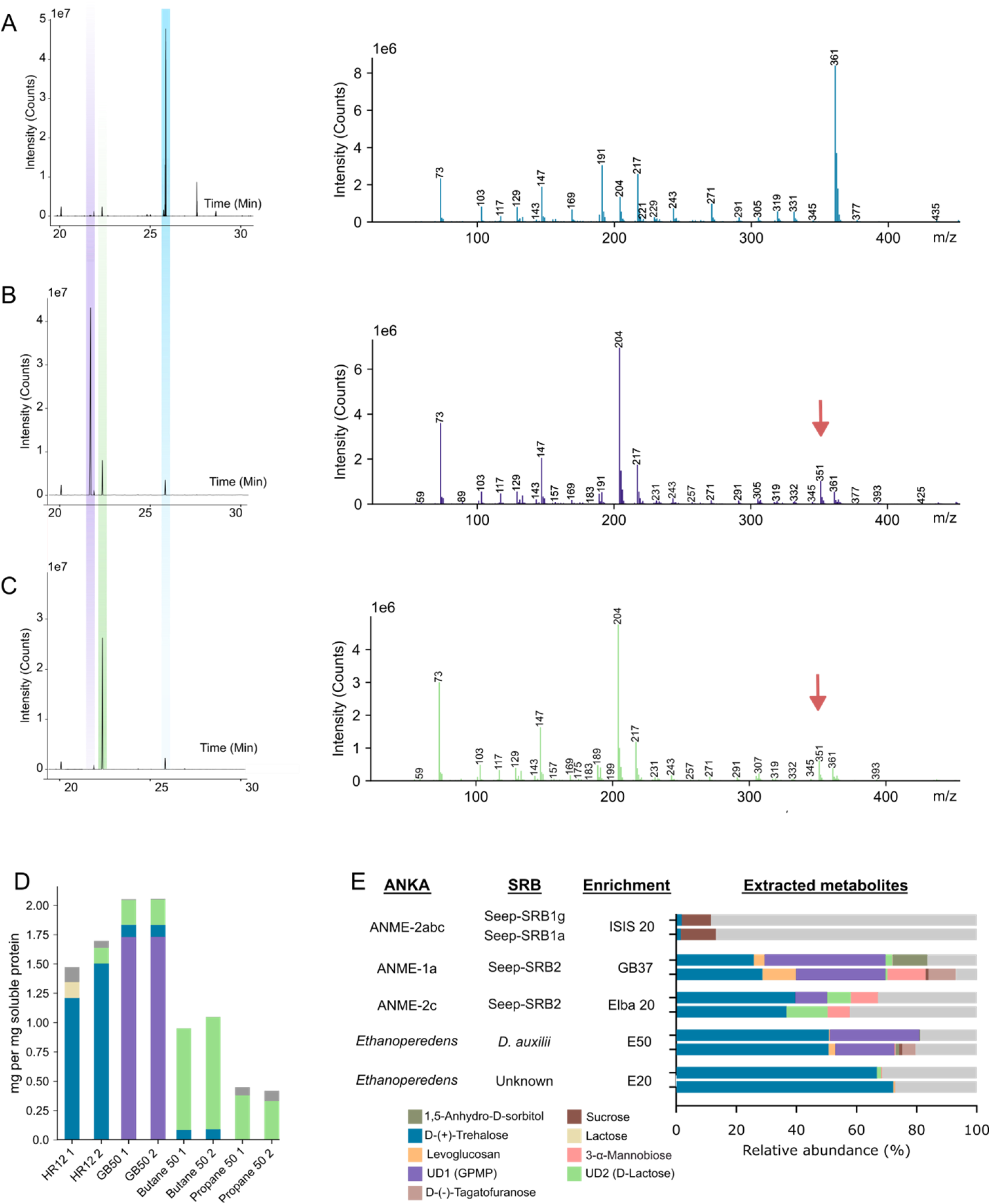
Extracted disaccharides and their mass spectra in alkane-oxidizing cultures. **A-C**) 20-30 min section of the chromatogram of the HR12, GB50, and Butane50 cultures and mass spectra of their most abundant disaccharide **A**) blue: trehalose at RT 25.86 min, **B**) purple: unidentified disaccharide isomer 1 at RT 21.70 min, purple arrow **C**) green arrow: unidentified disaccharide isomer 2 (RT 22.36). Compound identification was performed using the NIST Mass Finder 1.1 software, with match factors of 97 and 87 for trehalose and lactose, respectively. Each sugar is annotated with an octakis (trimethylsilyl), which has been removed for clarity. **D**) Disaccharide relative to soluble protein concentrations in select alkanotrophic enrichment cultures. **E**) Relative concentrations of extracted polar metabolites. The bar chart is labeled with the host enrichments and their constituent ANME/SRB consortia members. Highlighted in colors are sugars of particular interest as annotated by NIST, with the -silyl prefixes removed for clarity.

Abundant peaks were annotated using the NIST Standard Reference Database (NIST11) (Stein et al., 1987) and, if possible, identified by comparing the chromatographic retention times with those of standards. In the HR12 culture, the main disaccharide was identified as trehalose, based on retention time and spectral matching (i.e., m/z 361 as a terminal silylated hexose unit [C_15_H_33_O_4_Si_3_]^+^, m/z 204 and 217 for silylated pyranose ring, and 191 for silylated hexoses; Figure 1A; Medeiros and Simoneit, 2007). Trehalose reached a concentration of 1.5 to 2 mg per mg of soluble protein as assessed by microBCA assay (Supplementary Table 3-4). In contrast, the ANME-1-dominated GB50 culture contained only 0.2 mg trehalose per mg of soluble protein (Figure 1D). Instead, this metabolite extract contained large amounts of an as-yet-unidentified disaccharide, which elutes at 21.7 minutes (Figure 1B, 1D) and produces fragments characteristic of disaccharides, including m/z 361, 217, 204, 147, and 73. However, the glucose-associated m/z 361 signal had lower intensity, and the m/z 204 signal had higher intensity than those observed in the trehalose mass spectrum. In addition, we observed an abundant m/z 351 signal but could not link it to any known fragment. NIST annotates this compound as the lactose conformer 4-O-beta-Galactopyranosyl-D-mannopyranose (GPMG), but with a match factor <90. The Butane50 culture showed a peak at 22.4 minutes with fragmentation similar to that of the GB50 culture (Figures B-C, red arrow). Hence, the disaccharide peaks observed in the GB50 and Butane50 cultures may be isomers of a previously unreported disaccharide. To verify that this disaccharide was not GPMG, its elution was compared to a standard, which eluted much later at around 25.7 minutes and without the mass fragment m/z 351 (Supplementary Figure 1)

Next, we tested additional AOM cultures available in our laboratory for the presence of disaccharides. From these samples, only relative metabolite abundances could be determined. (Figure 1 E, Supplementary Table 1). All low-temperature AOM cultures (Elba 20, HR12, GB37) contained trehalose, whereas isomers of the GPMG-like disaccharide were more prominent in the high-temperature AOM enrichment GB50. Notably, the Isis20 culture was unique in that sucrose was the dominant disaccharide (Figure 1E). This is the only enrichment culture dominated by ANME-2a/2b and Seep-SRB1 partners (Schreiber et al. 2010). The low relative abundance and sucrose-dominated disaccharide composition of the Isis20 enrichment indicate that this consortium may exhibit distinct dynamics in modulating resource allocation within the consortium.

### 3.3 ANME and their partner bacteria encode disaccharide metabolism

To assess the potential of ANKA and their syntrophic SRB partners to produce and metabolize disaccharides, we constructed a genomic database including all available ANKA genomes (excluding ANME-3 lineages), genomes of characterized SRB partners from GTDB, and a representative subset of other Halobacteriota and Desulfobacterota genomes (Methods; Supplementary Table 6). We analyzed this database using hidden Markov models (HMMs) targeting key protein families involved in trehalose/sucrose metabolism and polysaccharide degradation (Table 2).

**Table 2:**
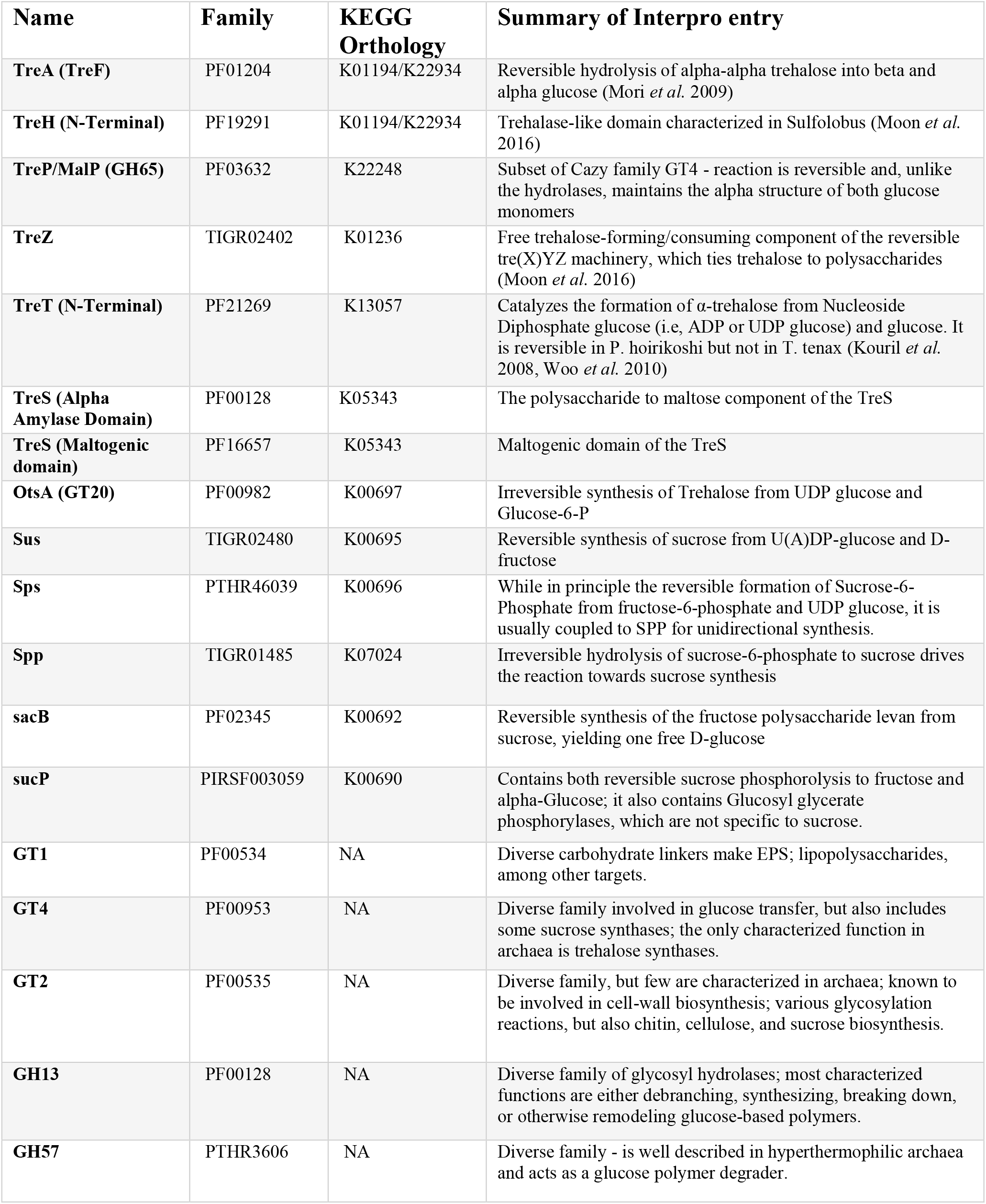
Protein family HMMs used in this study.

| Name | Family | KEGG Orthology | Summary of Interpro entry |
| --- | --- | --- | --- |
| **TreA (TreF)** | PF01204 | K01194/K22934 | Reversible hydrolysis of alpha-alpha trehalose into beta and alpha glucose (Mori *et al.* 2009) |
| **TreH (N-Terminal)** | PF19291 | K01194/K22934 | Trehalase-like domain characterized in Sulfolobus (Moon *et al.* 2016) |
| **TreP/MalP (GH65)** | PF03632 | K22248 | Subset of Cazy family GT4 - reaction is reversible and, unlike the hydrolases, maintains the alpha structure of both glucose monomers |
| **TreZ** | TIGR02402 | K01236 | Free trehalose-forming/consuming component of the reversible tre(X)YZ machinery, which ties trehalose to polysaccharides (Moon *et al.* 2016) |
| **TreT (N-Terminal)** | PF21269 | K13057 | Catalyzes the formation of α-trehalose from Nucleoside Diphosphate glucose (i.e, ADP or UDP glucose) and glucose. It is reversible in *P. hoirikoshi* but not in *T. tenax* (Kouril *et al.* 2008, Woo *et al.* 2010) |
| **TreS (Alpha Amylase Domain)** | PF00128 | K05343 | The polysaccharide to maltose component of the TreS |
| **TreS (Maltogenic domain)** | PF16657 | K05343 | Maltogenic domain of the TreS |
| **OtsA (GT20)** | PF00982 | K00697 | Irreversible synthesis of Trehalose from UDP glucose and Glucose-6-P |
| **Sus** | TIGR02480 | K00695 | Reversible synthesis of sucrose from U(A)DP-glucose and D-fructose |
| **Sps** | PTHR46039 | K00696 | While in principle the reversible formation of Sucrose-6-Phosphate from fructose-6-phosphate and UDP glucose, it is usually coupled to SPP for unidirectional synthesis. |
| **Spp** | TIGR01485 | K07024 | Irreversible hydrolysis of sucrose-6-phosphate to sucrose drives the reaction towards sucrose synthesis |
| **sacB** | PF02345 | K00692 | Reversible synthesis of the fructose polysaccharide levan from sucrose, yielding one free D-glucose |
| **sucP** | PIRSF003059 | K00690 | Contains both reversible sucrose phosphorylase to fructose and alpha-Glucose; it also contains Glucosyl glycerate phosphorylases, which are not specific to sucrose. |
| **GT1** | PF00534 | NA | Diverse carbohydrate linkers make EPS; lipopolysaccharides, among other targets. |
| **GT4** | PF00953 | NA | Diverse family involved in glucose transfer, but also includes some sucrose synthases; the only characterized function in archaea is trehalose synthases. |
| **GT2** | PF00535 | NA | Diverse family, but few are characterized in archaea; known to be involved in cell-wall biosynthesis; various glycosylation reactions, but also chitin, cellulose, and sucrose biosynthesis. |
| **GH13** | PF00128 | NA | Diverse family of glycosyl hydrolases; most characterized functions are either debranching, synthesizing, breaking down, or otherwise remodeling glucose-based polymers. |
| **GH57** | PTHR3606 | NA | Diverse family - is well described in hyperthermophilic archaea and acts as a glucose polymer degrader. |

Among the ANKA lineages, *Ca*. Ethanoperedens genomes (n = 15) encode the greatest functional potential for disaccharide metabolism, with an average of 6.1 genes per genome matching the tested HMM profiles (Figure 2). *Ca*. Alkanophagaceae (4.7 genes per genome, n = 3) and ANME-2c (4.5 genes per genome, n = 22) also generally encode multiple key enzymes, including trehalose synthase (TreT), trehalose-maltose phosphorylase (TreP/MalP), and trehalose-6-phosphate synthase (OtsA). These values are typical for Halobacteriota (4.1, n=289). Other ANME groups, including ANME-2a/2b (2.6, n = 42), ANME-2d (3.3, n = 54), and ANME-1 (2.9, n = 87), have, on average, far fewer genes encoding disaccharide metabolism. Notably, variants that catalyze the irreversible degradation of trehalose are absent in all ANKA. *Ca*. Methanoliparaceae had the fewest genes encoding disaccharide metabolism (2.3 hits per genome). Hence, both *Ca*. Methanoliparaceae and the ANME-2d may not be able to metabolize trehalose or other disaccharides (Figure 2).

**Figure 2.**
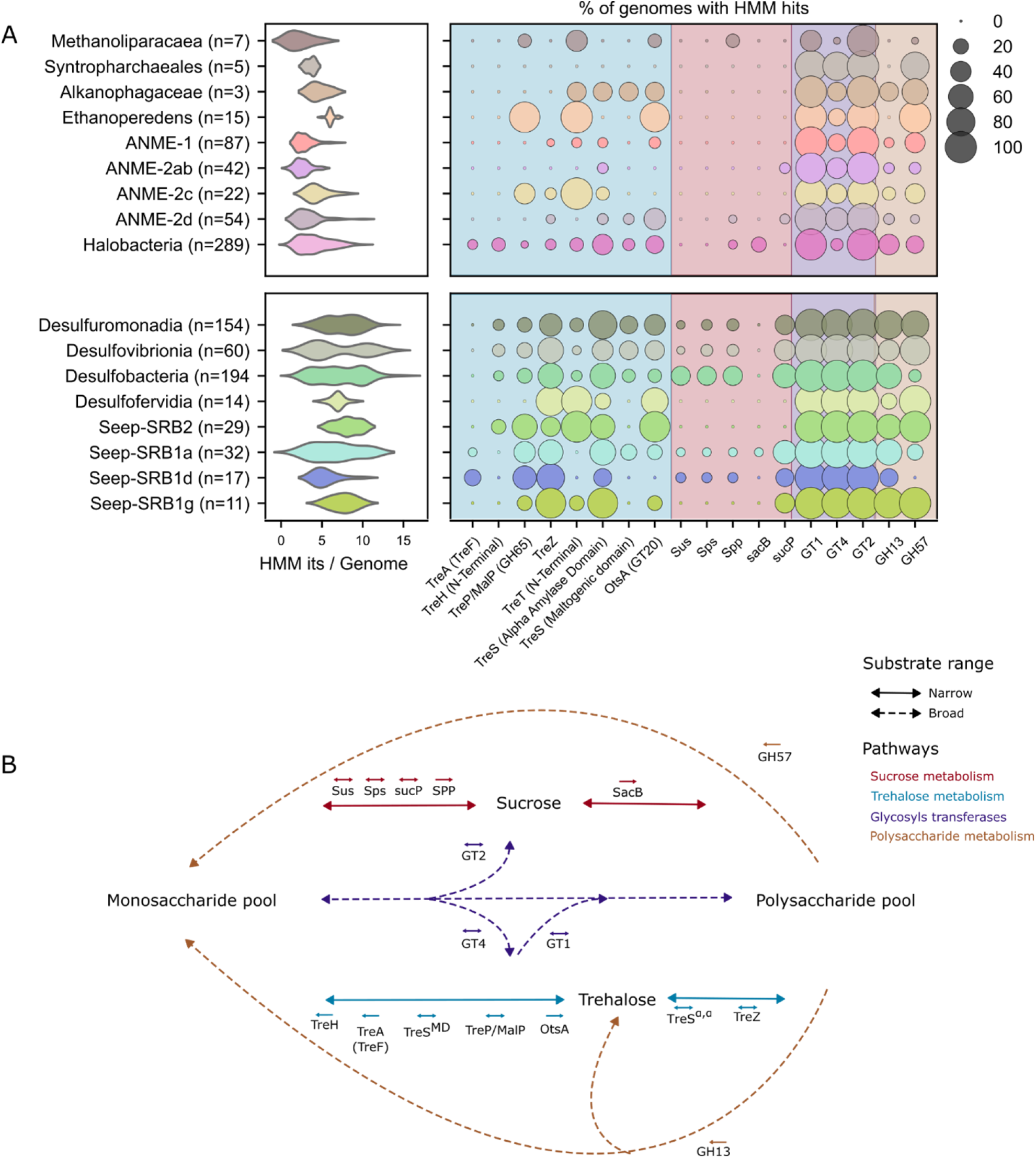
Genes encoding sugar metabolism in the genomes of ANKA, partner SRBs, and reference organisms. **A**) Presence of genes encoding trehalose-, sucrose-, and related carbohydrate metabolism across all GTDB (R220)–deposited genomes of ANKA and known SRB partners, as well as selected Desulfobacterota and non-ANKA Halobacteriales genomes according to HMM profiles. Hits were retained for e-values ≤ 1e-10 and HMM profile coverage ≥ 40%. Each gene was assigned to only one HMM profile. The left panel shows the number of hits per genome for each taxonomic group; the central panel shows the percentage of genomes in each group that contain at least one hit associated with the queried HMM profiles. Blue indicates reactions directly operating on trehalose, red indicates sucrose, and purple indicates glycosyl transferase families that may operate on trehalose or sucrose. In contrast, gold indicates select glycosyl hydrolases that degrade polysaccharides. **B**) Schematic overview of known enzymes for disaccharides trehalose (blue) and sucrose (red) metabolisms as well as select polysaccharide degradation (gold) and glycosylation metabolisms (purple) that intersect with the carbohydrate pools investigated in this study. Enzymes with specific substrates are shown with solid lines, while enzymes with broad substrate ranges are shown with dashed lines. Above each enzyme name is an arrow which indicate its known directionality. The enzymes used for HMM profiles are listed in Table 2. Abbreviations: TreA (TreF) – trehalase; TreH – reversible α,α-trehalose hydrolase; TreP/MalP – trehalose/maltose phosphorylase; TreZ – trehalose-6-phosphate hydrolase; TreT – trehalose glycosyltransferase; TreS – maltose-forming trehalose isomerase; OtsA – trehalose-6-phosphate synthase; Sus – sucrose synthase; Sps – sucrose-phosphate synthase; Spp – sucrose-phosphate phosphatase; SacB – levansucrase; SucP – sucrose/glucosylglycerate phosphorylase; and broad glycosyltransferase (GT1, GT2, GT4) and glycoside hydrolase (GH13, GH57) families.

In contrast, members of the Desulfobacterota exhibit far greater functional diversity in the distribution of disaccharide metabolism-related genes. The average number of genes with HMM-profile hits per genome ranged from 5.5 in Seep-SRB1d to 8.1 in Seep-SRB2. All SRB clades known to engage in syntrophic partnerships with ANKA, such as Seep-SRB1a, SRB1g, SRB2, and *Ca*. Desulfofervidus auxilii encode trehalases such as TreH and TreA, which catalyze the irreversible degradation of trehalose, and trehalose synthases, including TreS and TreT, which catalyze the formation of trehalose. Seep-SRB1a most reflected the distribution of Desulfobacteria, while other classes of the Desulfobacterota, like Desulfovibrionia and Desulfuromonadia, display bimodal distributions of HMM hits per genome, indicating that there is likely sub-class level differentiation in disaccharide metabolism. (Figure 2). While not common in the SRB, TreH is sporadically present in the Seep-SRB2 and the classes that do not contain known ANKA partners (Figure 2), while TreA is common in the Seep-SRB1d and rare or absent in the other SRB. In contrast to ANKA, many SRB partners encode trehalase, suggesting that only these members can break down trehalose. The presence of genes responsible for the reversible synthesis of trehalose in ANKA and irreversible trehalose degradation in SRB suggests a potential link in trehalose metabolism between ANKA and their partners.

### 3.4 ANKA and their partner bacteria share trehalose metabolism via horizontal gene transfer

To elucidate the evolutionary history of trehalose metabolism in ANKA and SRB partners, we reconstructed amino-acid sequence-based phylogenies for three key genes involved in trehalose biosynthesis and/or degradation: the trehalose-6-phosphate synthase (OtsA), the trehalose/maltose phosphorylase (TreP/MalP), and the trehalose glycosyl-transferring synthase (TreT). These enzymes represent the most common enzymes involved in trehalose synthesis, breakdown, and (de)polymerization across both SRB and ANKA (Figure 2A), and their distribution between the ANKA-SRB functional groups offers valuable insights into patterns of vertical inheritance and horizontal gene transfer (HGT) events.

Trehalose/maltose phosphorylase (TreP/MalP) catalyzes the reversible breakdown of trehalose or maltose into glucose (Table 2, Figure 2B) and is common in *Ca*. Ethanoperedens and ANME-2c, but appears only in one of seven *Ca*. Methanoliparacea. While TreP/MalP occurs in <20% of the ANKA genomes, it is present in all SRB partner clades except *Ca*. Desulfofervidia. This particular protein family contains both trehalose- and maltose-metabolizing members, and the majority of the hits retrieved are more homologous to a characterized trehalose phosphatase than to a characterized maltose phosphatase (Figure 3A). The TreP/MalP coding sequences of ANME-2c form a separate phylogenetic clade with a single member of the *Ca*. Ethanoperedens, indicating an HGT between these two ANKA (Figure 3A). The TreP sequences of the remaining *Ca*. Ethanoperedens genomes form two distinct phylogenetic clusters. EX4572-44 including *Ca*. Ethanoperedens thermophilum, likely acquired *treP/malP* from a Syntrophotalea-like ancestor, while *Ca*. Ethanoperedens ethanivorans (originally described as *Ca*. Argoarchaeum ethanivorans; Chen et al. 2019) may have acquired *treP* from a Seep-SRB2-like ancestor. This indicates that the ancestors of psychro- and thermophilic extant ethane oxidizers acquired *treP* independently of one another. Similarly, the TreP-encoding sequences of cold-growing Seep-SRB1a are separated into two phylogenetic clades: one is phylogenetically distinct, and the other is shared with members of Seep-SRB1g and SRB1d and appears to be rooted in a shared ancestor with Desulfovibrio.

**Figure 3.**
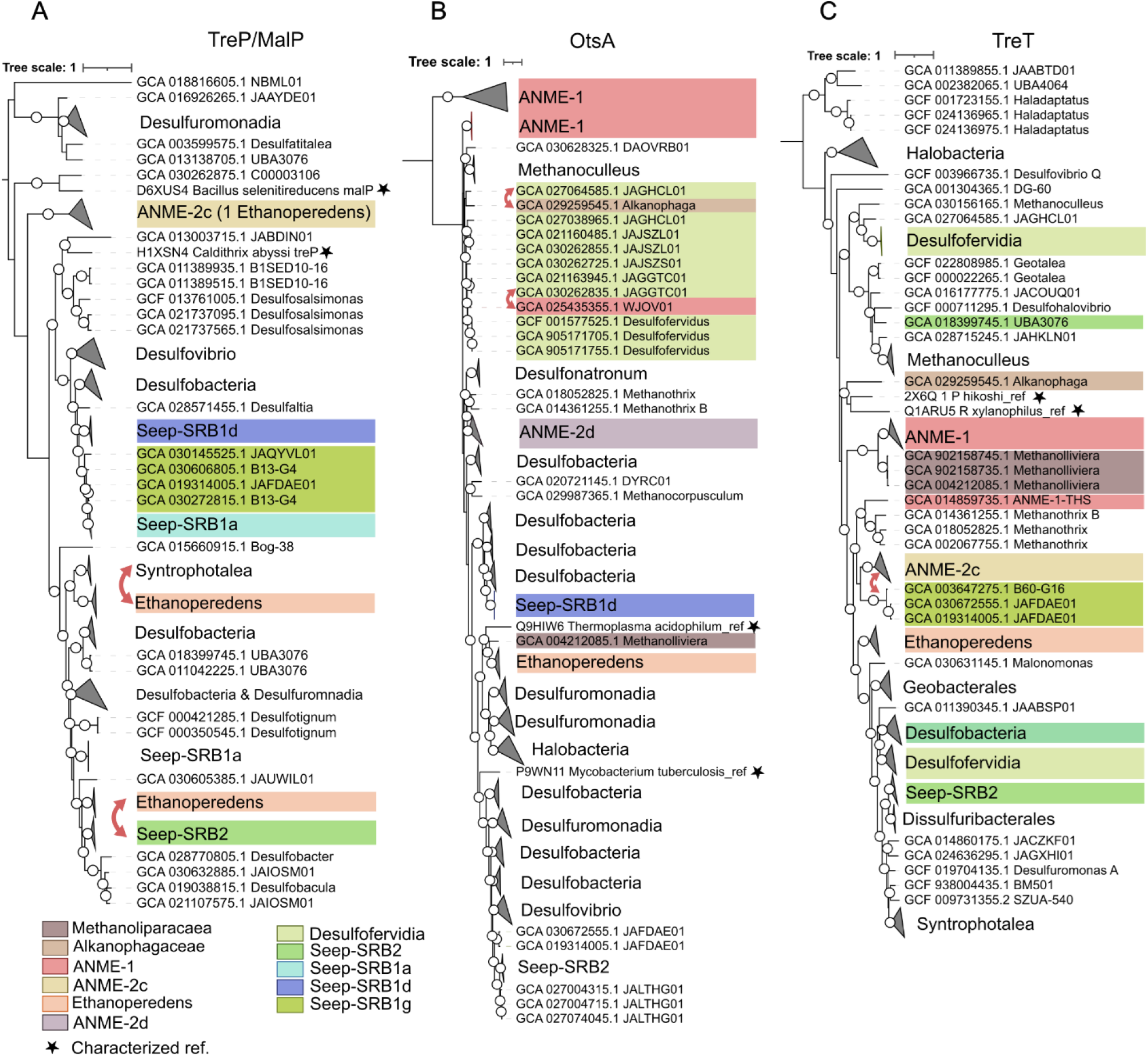
Horizontal gene transfer (HGT) between ANKA and SRB partners of the trehalose-metabolizing enzymes OtsA, TreP/MalP, and TreT. Protein trees of best-hit locus tags to the HMMs corresponding to the protein families of **A**) TreP/MalP (GH65), **B**) OtsA (GT20), and **C**) TreT. The branches are shaded according to their membership in an ANKA or SRB partner functional group, as in Figure 2, and proteins with characterized references are shown in purple. Red arrows mark potential ANKA/SRB HGT events. In all trees, bootstraps greater than 90% are shown as white circles. Grey stars show characterized references, and functional and taxonomic groupings of interest are highlighted by boxes following the color scheme of Figure 2.

OtsA, the trehalose-6-phosphate synthase, catalyzes the irreversible conversion of UDP-glucose to trehalose and is found in representatives of all three domains of life (Avonce et al., 2006). In our metagenomic dataset, sequences encoding OtsA separate into two distinct clades, one of which is composed exclusively of ANME1 (Figure 3B). One of these clades is a cluster of four proteins, all annotated as D-inositol-3-phosphate glycosyltransferases and represent the lower limit of our similarity selection criteria (Supplementary Table 7). Beyond these, only 6 of the 95 Syntropharchaeales genomes contain an OtsA homolog. Two of these are clear cases of HGT for ANME-1 and *Ca*. Alkanophaga genomes (Figure 3B), as these two instances of this gene in these two archaea exist in isolation inside an otherwise bacterial clade. In both cases, the closest OtsA homologs derive from their SRB partners, the thermophilic *Ca*. Desulfofervidia. On the other hand, ∼ 45% of ANME-2d genomes and all but two *Ca*. Ethanoperedens genomes encode OtsA homologs (Figure 2B, Supplementary Table 6). However, the phylogenetic reconstruction of OtsA reveals that these closely related ANKA groups have acquired *otsA* independently (Figure 3A). The ANME-2d sequences form a clade with Methanothrix, consistent with vertical inheritance from a shared ancestor. *Ca*. Ethanoperedens and one MAG of *Ca*. Methanolliviera seemed to have acquired *otsA* from a common ancestor with the Desulfuromonadia. *otsA* appears to have been acquired twice independently within the Desulfuromonadales, where one clade of OtsA shares an ancestor with the *Ca*. Ethanoperedens and the other clade appear to preserve bacterial phylogeny due to a shared root with the characterized Mycobacterium reference. In contrast, a second Desulfuromonadales group is nested within the Desulfobacterales in the OtsA tree, suggesting HGT from within that order. This latter group includes the acetylene-degrading Syntrophotalea, which appears to have acquired *otsA* horizontally from a Desulfobacterales source.

TreT, which catalyzes the reversible formation of trehalose from glucose (Table 2), also shows signs of interdomain HGT. Homologs of this enzyme are present in most ANME-2c and *Ca*. Ethanoperedens genomes, with sporadic occurrences in the Methanliparacaea and *Ca*. Alkanophaga. TreT is also found in Seep-SRB2 and Desulfofervidia, but is less common in free-living Desulfobacterota. TreT sequences encoded by Desulfofervidia form two different phylogenetic clades (Figure 3C). One of these clades appears to root with a shared ancestor with the hydrogenotrophic Methanoculleus, indicating HGT between SRB and non-ANKA members of the Archaea. The TreT encoding sequences of other Desulfovibrio are closely related to TreT proteins of the Desulfobacteria, including syntrophic members like the Seep-SRB2, and members of the Desulfuromonadia, including Syntrophotalea. Moreover, this TreT clade shares an ancestor with the Ethanoperedens and other ANKA. OtsA and TreT/MalP of the Halobacterota and Desulfobacterota appear to be rooted in anaerobes, but TreT appears to be ancestral to the aerobic and halophilic archaea of the Halobacteria.

### 3.5 ^13^C trehalose and ^13^C-DIC assimilation reveal active, alkane-dependent heterotrophs

We hypothesized that small disaccharides such as trehalose may play a role in the syntrophic relationship among ANKA, SRB, and ancillary community heterotrophs. To probe the role of disaccharides in ANKA/SRB enrichment cultures, we performed two ^13^C-labeling experiments with the thermophilic Ethane50 culture. In the first SIP experiment, we sought to investigate the role of disaccharides as potential carbon carriers for alkane-oxidizing consortia and, eventually, for use by ancillary community heterotrophs. Trehalose was selected due to its high concentration in the Ethane50 culture (Figure 1) and the abundant enzymatic machinery of both the constituent ANKA (*Ca*. E. thermophilum) and SRB (*Ca*. D. auxilii) for trehalose metabolism (Figure 2). Aliquots of the culture were incubated with *Ca*. 10% ^13^C-labeled trehalose, either with or without unlabeled ethane, and sampled over 15 days (Figure 4A). Incubations of Ethane50 with trehalose showed notable sulfide formation and an increase in δ^13^C values of DIC regardless of the presence of ethane (Supplementary Tables 9 and 10, Supplementary Figures 2-3), indicating that organoheterotrophs degrade the majority of supplied trehalose within one week (δ^13^C-values w/o ethane = 279 ‰, with ethane = 287 ‰, Supplementary Figures 2-3, Supplementary Table 10). The ^13^C incorporation into lipids was tracked to identify the actors in trehalose utilization and immediate δ^13^C-values (>1000‰ within hours) and highest (> 3000‰ within <7 days) were obtained for the terminally-branched fatty acids iC15:0, iC16:0, and iC17:0, recently identified to be associated with heterotrophic organisms (Zhu et al., 2022). The rapid and intense incorporation is notable, given that the culture had not previously been supplemented with external trehalose, and indicates that the heterotrophs are well adapted to respond quickly to changes in disaccharide availability, using trehalose as both a carbon and an energy source. In contrast, other fatty acids primarily produced by the SRB partner bacterium (Zhu et al., 2022) assimilated much less trehalose (δ^13^C16:0 < 400‰; and C18:0 < 30‰) (Figure 4A). The C16:0 fatty acid is also potentially produced in part by heterotrophs; hence, the higher δ^13^C values could be attributed to heterotrophic production (Heinzelmann et al., 2015). The small changes in δ^13^C value for C18:0 may also be a result of secondary DIC uptake, as ^13^C-labeled carbon is readily transferred into the DIC pool. The archaeal markers of phytane (δ^13^C <10‰) and biphytane (δ^13^C <−40‰) showed minor ^13^C incorporation, although an order of magnitude below the transfer to the DIC pool; however, the incorporation was independent of the presence of ethane. This implies that the ethane-oxidizing archaea are heterotrophically assimilating carbohydrates, as suggested by Yin et al. (2022). Alternatively, this could be explained by heterotrophic archaea, which can produce these isoprenoids (He et al., 2022) and are present in low abundances in these enrichments (Hahn et al., 2020).

**Figure 4.**
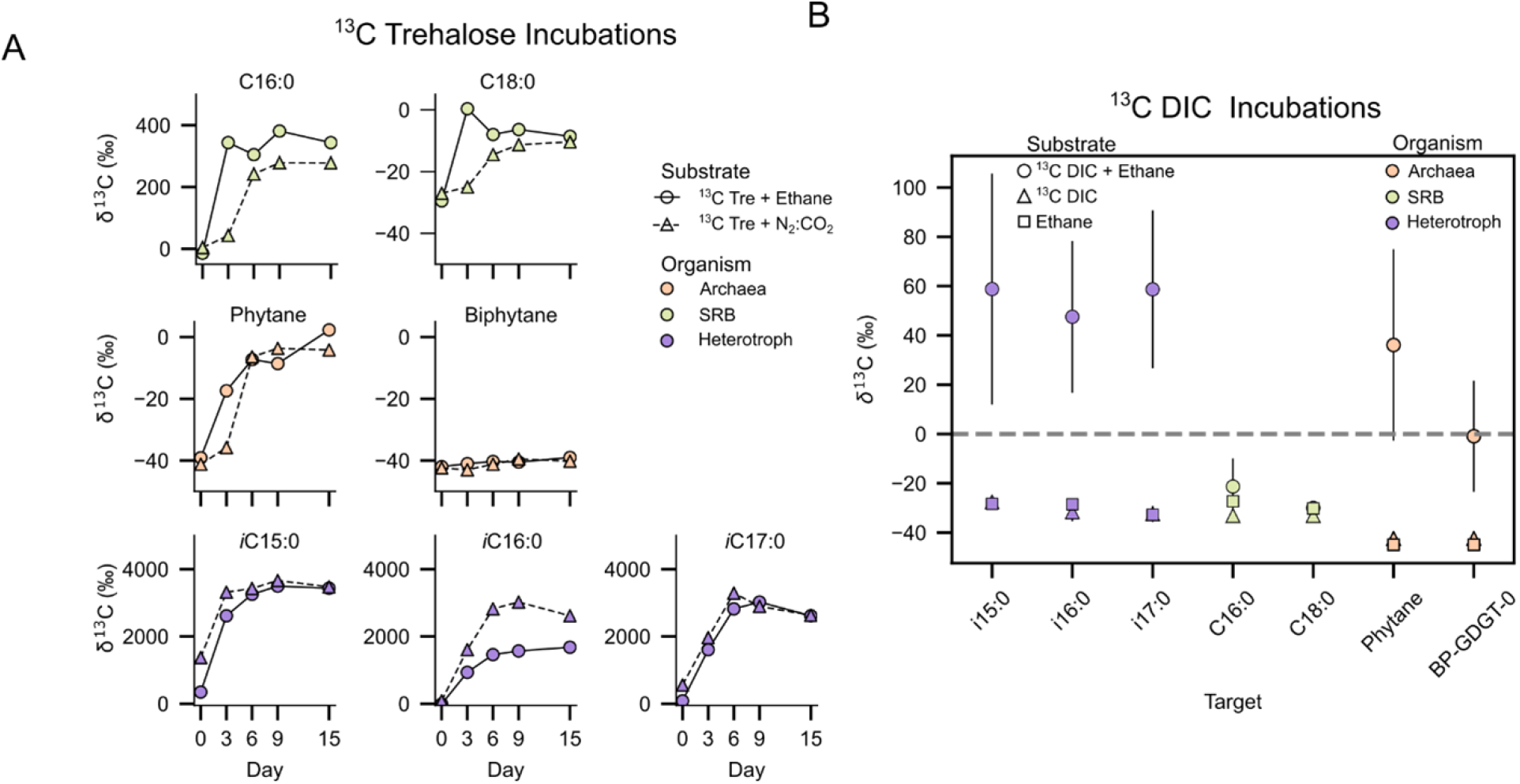
Incorporation of ^13^C-trehalose and ^13^C-DIC into bacterial and archaeal lipid moieties, in the Ethane50 culture with and without ethane. **A)** Shift in 13C values of marker lipids signaling heterotrophs (purple, 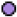), SRB (light-green, 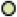), and ANKA (light-orange, 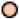) during trehalose incorporation over 15 days. Data points represent measurements from single incubations. Here, biomass constraints forced a trade-off between replicates and time-series resolution, so there are no replicates. Biomass was distributed into separate Schott flasks, with one bottle for each condition and time point. These bottles were then sampled in their entirety at the corresponding time point for the analysis of δ^13^C incorporation into lipids (Supplementary Table 11). Sulfide was used to track activity (Supplementary Tables 8 and 9). **B**) Change of δ^13^C values of the same lipids during DIC incorporation after 24 days. Here, each data point is the mean of triplicate incubations with standard deviation visualized as error bars (Supplementary Table 12). Activity was tracked by sulfide formation (Supplementary Table 9). Abbreviation: BP-GDGT-0, Biphytane-glycerol dialkyl glycerol tetraether without rings.

To resolve the ethane dependency of heterotrophs on the activity of ANKA-SRB consortia, we incubated the Ethane50 culture with ^13^C-DIC, either with or without ethane as the only energy source (Figure 4B, Supplementary Fig. 5-7). As expected, the incorporation of DIC into the isoprenoids produced by *Ca*. Ethanoperedens fully depended on the presence of ethane (δ^13^C phytane=36 ± 39‰; biphytane = −1 ± 22‰) after 24 days of incubation (Figure 4A). Likewise, ^13^C-DIC assimilation into the fatty acids predominantly produced by the partner SRB depended on the presence of ethane, although these lipids show only limited or nominal incorporation from DIC (δ^13^C-values C16:0 = 11 ± 11‰, C18:0 = 18 ± 22‰). On the other hand, DIC-derived carbon was substantially incorporated into the fatty acids that are typically produced by ancillary heterotrophic bacteria (δ^13^C-values for iC:15:0 = 59 ± 47‰, iC:16:0 = 48 ± 30‰, iC17:0 = 59 ± 30‰). This experiment demonstrates that both the *Ca*. Ethanoperedens/SRB core community and the associated heterotrophic bacteria depend on the energy of ethane oxidation when no exogenous carbohydrates are supplied.

The trehalose incubation experiments suggest that neither *Ca*. E. thermophilum nor the partner bacterium *Ca*. D. auxilii assimilated substantial amounts of externally provided trehalose, but ancillary community heterotrophs readily consumed that trehalose. The DIC incorporation experiment shows that these community heterotrophs only assimilate DIC-derived carbon when ethane is present. Considered together, these experiments demonstrate that ancillary community heterotrophs likely receive carbohydrates derived from DIC carbon synthesized by the ANKA/SRB consortia.

## 4 Discussion

This study demonstrates the accumulation of substantial amounts of trehalose and other disaccharides in sediment-free, alkane-degrading cultures consisting of ANKA and their bacterial partners. In particular, in the cold-adapted methane-oxidizing and cold- and hot-adapted ethane-oxidizing cultures, trehalose is abundant, reaching 1.5–2 mg trehalose per mg soluble protein (Figure 1). This is almost twice the highest levels reported so far for the archaeon *Haladaptus paucihalophilus* when cultured under extreme osmotic stress (Youssef et al., 2013), but within the highest ranges reported for halophilic SRB responding to salt stress (Welsh et al., 1996). In other cultures, such as GB50 or Butane50 from the Guaymas Basin, other, yet-unidentified disaccharides are more abundant. Given that alkanes and sulfate serve as primary energy sources and electron sinks, respectively, and that ANKA and their partner bacteria strongly dominate the microbial communities, it is highly likely that these disaccharides are metabolic products of anaerobic alkane oxidation.

Our results suggest that ANKA, rather than partner bacteria, produced most of the disaccharides in alkane-oxidizing cultures. This follows from the fact that AOM50, Ethane50, and Butane50 contain specific ANKA lineages but share *Ca*. Desulfofervidus as a partner bacterium (Laso-Pérez et al. 2016, Hahn et al. 2020)(Figure 5). Despite sharing bacterium, these cultures exhibit markedly different disaccharide profiles, suggesting that the archaeal partner exerts primary control over the disaccharide pool.

**Figure 5.**
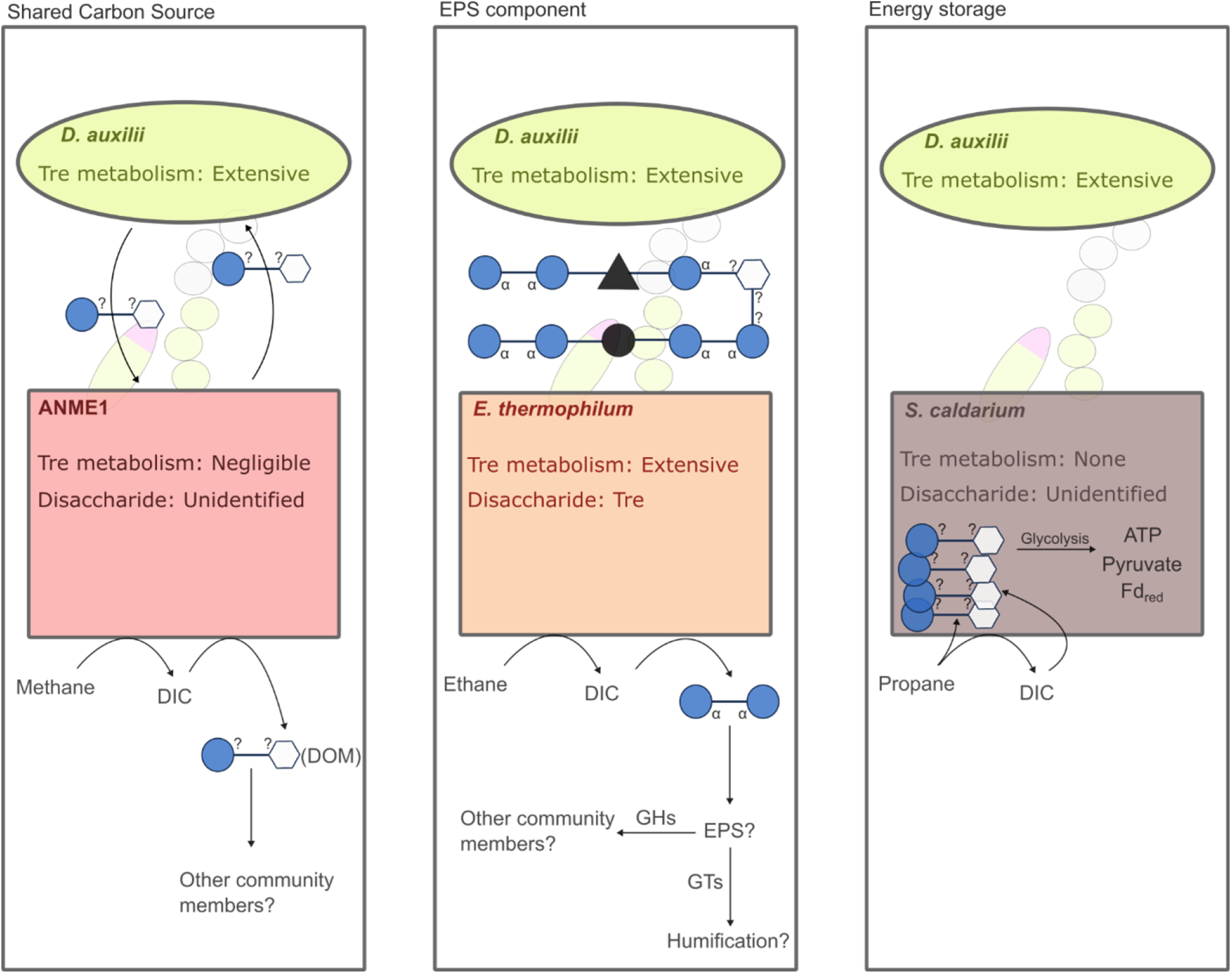
Potential roles of disaccharides in ANKA beyond compatible solute. Depicted are illustrations of three ANKA enrichments that share the same SRB partner with the archaea oxidizing methane (left), ethane (middle), and propane (right). All of these were shown to have substantial concentrations of disaccharides. Of these enrichments, only the Ethane50 enrichment containing *Ca*. E. thermophilum, which encodes extensive trehalose metabolism, was found to have trehalose as its main disaccharide, with unidentified disaccharides in the enrichments of ANME1 and Syntropharchaeum, and the metabolism is only shown for the archaeal partner. Note that this figure is illustrative and does not imply any level of certainty regarding which partner synthesized the disaccharide. Glycans are shown according to the SIGAF nomenclature, with black circles and triangles representing non-carbohydrate components such as quinones, amino acids, or hemes. Abbreviations: DIC, dissolved inorganic carbon; DOM, dissolved organic carbon; EPS, Extracellular Polysaccharide; GHs, Glycosyl Hydrolases; GTs, Glycosyl Transferases.

However, the partner bacteria may also contribute to the disaccharide pool. In trehalose-producing cultures, trehalose metabolism was present in both archaea and partners. The unidirectional trehalose-degrading enzymes appear completely absent from all ANKA genomes and occur only sporadically in partner bacterial groups. Reversible trehalose degradation or synthesis enzymes are common in ANKA from all trehalose-containing cultures and across all partner bacteria groups. Trehalose synthesis in SRB has been suggested based on metagenomic data (Murphy et al., 2021; Spring et al., 2019), but has only been shown biochemically in Desulfovibrio halophilus, where it increases under salt stress (Welsh et al., 1996). Similar salt-stress-dependent trehalose production has been shown for Halobacteria (Youssef et al. 2013). In our recent study, we show that the isotope values of disaccharides in AOM-active sediments are more similar to those of lipids associated with SRB (Stock et al., 2026). However, *Methanosarcina barkerii*, a close relative of the *Ca*. Ethanoperedens and the ANME2 have been reported to exhibit substantially more negative δ13C values in lipids than in bulk biomass, particularly under substrate-limiting conditions (Londry et al., 2008). Similarly, *in situ* AOM isotope fractionation is likely also constrained by variable substrate availability and the rates of syntrophic electron transfer, as implied by decades of findings in both enrichment and environmental studies (McGlynn et al., 2015; Nauhaus et al., 2005; Timmers et al., 2015; Wegener et al., 2022, 2016). Thus, the fact that the isotope values of the sugars we report in Stock et al. (2026) are less depleted than the archaeal lipids is not conclusive evidence that SRB produce the disaccharides; the origin of these deep-sediment sugars remains an open question requiring biochemical verification.

### 4.1 Role of disaccharides in alkane-oxidizing cultures

The functional role of the disaccharides in alkane-oxidizing cultures remains an intriguing unknown. The ANKA/SRB consortia may produce disaccharides as a catabolic energy storage mechanism. Energy storage might be essential for highly specialized, presumed immobile organisms that can only grow in the narrow, fluctuating sulfate-methane (or alkane) interface. For SRB, an energy storage in disaccharides would be notable, because SRB typically favor glycogen (Stams et al., 1983) or polyhydroxyalkanoates (PHAs) as storage compounds (Hai et al., 2004). However, trehalose or other sugars could conceivably be converted into these compounds via known pathways (Figure 2), or ANKA may use disaccharides as an energy reserve during alkane and/or sulfate limitation. In the presence of sulfate, the disaccharides could be metabolized glycolytically into acetate, which could then be oxidized to CO_2_, while reducing equivalents can be transferred to the partner bacterium. In the absence of sulfate, ANKA could potentially ferment the disaccharide and emit compounds such as lactate or acetate. Future culture starvation experiments should test whether AOM biomass releases sugar-derived CO_2_ or fermentation products when starved of electron acceptors or donors.

Trehalose is a well-known osmolyte in marine organisms (Gregory and Boyd, 2021; McParland et al., 2021; Poli et al., 2017). However, our alkane-oxidizing cultures were maintained at a constant salinity of 35‰, which closely mimics the organisms’ native conditions, making osmotic stress unlikely to drive these extreme accumulations of trehalose or yet-unknown disaccharides. Instead, these disaccharides may serve as precursors for extracellular polymeric substances (EPS) (Figure 5). EPS is known to play a role in ANME consortia and has been suggested to participate in microbially induced silica precipitation (Osorio-Rodriguez et al., 2023). While the 1,1-glycosidic linkage of trehalose precludes linear polymerization, certain trehalose-maltose phosphorylases or maltose-forming trehalose isomerases can interconvert trehalose and maltose, facilitating the formation of 1,4-linked polymers (Koh et al., 2003). Trehalose could serve as a basis for other polymers linked by compounds such as proto-humics (quinones) (Figure 5). Such polymers have been synthesized by humans (Kukowka and Maślińska-Solich, 2010; Oliveri et al., 2016; Vinciguerra et al., 2022), but have not been reported from natural environments. Such “humified” sugar derivatives could provide a protective matrix for the consortia and an electrically conductive matrix to support DIET.

In addition, such a humified sugar matrix may explain the reports on the presence and formation of “amorphous carbon” in ANME archaea (Allen et al., 2021). This study extracted AOM cultures by sequential boiling in strong bases and acids. The residue of this procedure was described as amorphous carbon. However, the boiling of AOM biomass rich in polysaccharides, proto-humics like quinones, and Fe-S minerals, would promote sulfur incorporation and condensation reactions that yield sulfurized organic matter similar to humin (Raven et al., 2021; van Dongen et al., 2003). The subsequent acid boiling step could further complexate the base-insoluble biomass, as has been found for other types of organic matter (Bellera et al., 2015). Procedures very similar to those employed by Allen et al (2021) to extract amorphous carbon have been employed to transform algal biomass into recalcitrant humic-like compounds (Kok et al., 2000). The resulting base-insoluble, blackened biomass would produce Raman spectra similar to those previously reported for multiple humic substances (Sánchez-Cortés et al., 1998; Yang et al., 1994), with the additional caveat that Li et al. have recently reported that Raman spectroscopy can induce the D and G bands used to identify carbonaceous compounds in otherwise non-carbonaceous organic matter (Li et al., 2023). Together, these observations suggest an alternative, extraction-based origin of the refractory black organic matter in AOM cultures.

While the function of disaccharides in ANKA/SRB consortia remains unknown, their exudation appears to have substantial effects on the surrounding microbiome. Deep-sea sediments contain high numbers of heterotrophic microorganisms while being poor in potential substrates (Arnosti et al., 2019). Recent work suggests that some heterotrophs grow on AOM-derived proteinogenic necromass (Zhu et al., 2022). Trehalose and other disaccharides are also readily utilized as carbon sources by diverse heterotrophic populations (Lian et al., 2021). The presence of such heterotrophs in our enrichments (Laso-Pérez et al. 2016; Hahn et al. 2020), which immediately consume the provided trehalose, supports the idea that metabolites produced by ANKA-SRB consortia support parts of the surrounding heterotrophic sediment communities (Figure 5). The persistence of a large disaccharide pool despite active heterotrophs suggests an active sugar economy sustained by the ANKA–SRB consortia. The ANKA consortia may sustain a deep-sediment “barrier biome”, in which the oxidation of alkanes is coupled with the secretion of disaccharides that support a metabolically integrated assemblage of secondary heterotrophs. In this model, distinct ANKA–SRB pairings generate characteristic sugar signatures that shape the surrounding microbial community (Figure 5). What, if anything, the microbial community provides in return is unknown. Although these observations are derived from enrichment cultures, they point to a previously unrecognized in situ carbon exchange system. Deciphering this metabolic economy will refine our models of sedimentary carbon cycling and reveal how one of Earth’s largest microbial habitats manages its carbon flux.

## 5 Methods

### 5.1 Cultivation and maintenance of alkanotrophic enrichment cultures

The enrichment cultures used in this study are described in Table 1. Briefly, A collection of anaerobic enrichment cultures representative of alkane-oxidizing microbial consortia was produced from a range of geochemically and geographically distinct marine environments (Table 1). These cultures span a diversity of substrates (methane, propane, butane), archaeal lineages, incubation temperatures (12–60 °C), and original sediment depths (12–1999 m).

Low-temperature methane-fed enrichments included HR12 and Elba20, both of which were dominated by ANME-2c and Seep-SRB2. HR12 was derived from 780 m depth at the Cascadia Margin (SO148-1 expedition, 2000; Holler et al., 2009), while Elba20 was established from shallow sediments (12 m) near Elba Island (Hydra Diving Team, 2010; Krukenberg et al., 2018). A 20 °C enrichment from the ISIS mud volcano (RV L’Atalante, 2003) at 1250 m (ISIS20) included a mixed archaeal community comprising ANME-2c, ANME-2a, and ANME-2b with Seep-SRB1a (Schreiber et al. 2010)

Thermophilic cultures derived from hydrothermally active and hydrocarbon-containing Guaymas Basin sediments. The GB50 (50 °C) dominated by ANME-1a (AT15-45 expedition, 2009) included partnered with *Ca*. Desulfofervidus auxilia; (Krukenberg et al., 2018). Two additional 50 °C enrichments, fed with butane and propane, were dominated by *Ca*. Syntropharchaeum butanivorans and *Ca*. Syntropharchaeum caldarius, respectively, both paired with *Ca*. D. auxilii (Laso-Pérez et al. 2016). Two enrichments fed with ethane were investigated in this work, both from the AT37-06 expedition and enriched from the Guaymas Basin. Ethane50 was maintained at 50 °C (Hahn et al., 2020), and the other was maintained at 20 °C.

All cultures were produced and maintained as described by Laso-Pérez et al. (2018). Briefly, in an anaerobic chamber, the anoxic sediment was diluted with anoxic synthetic sulfate reducer medium (Widdel and Bak, 1992) at pH 7.2 (HR12, ISIS20) or at pH 6.5 for all other cultures. The resulting slurries were distributed into cultivation bottles, which, after the addition of gaseous substrates (methane, ethane, propane, or butane), were stored at the respective temperatures (Table 1). As a measure for substrate-dependent activity, sulfide concentrations were repeatedly measured in a photometric copper sulfate assay (Cord-Ruwisch, 1985). Unless otherwise specified, the sulfide was measured against a standard curve composed of dilutions of a 25 mM standard, the concentration of which was regularly checked by iodometric titration (American Public Health Association, 1980). The medium was exchanged when sulfide values approached or exceeded 15 mM. Successive dilutions of the slurries (1:2 to 1:4) led to virtually sediment-free enrichment cultures after 8 to 10 dilutions or 1-3 years.

### 5.2 Extraction of polar and non-polar metabolites

For sampling, the incubation bottles were inverted to allow the biomass to settle on or near the butyl rubber stopper. Then, 2 to 5 mL of the flock was sampled using plastic syringes with large-gauge (11) hypodermic needles into 15 mL centrifuge vials. The samples were centrifuged, and the supernatant was transferred into another Sarstedt tube. To extract polar metabolites from the biomass flocks, 2 mL of 1:1 water:MeOH was added. If samples were designated for quantification, the samples were spiked with ribitol (0.0714 mg mL^−1^) and cholestane (0.0286 mg mL^−1^). Samples were then sonicated using a Bandelin Sonoplus with an MS72 probe for 1 minute at 97% power. The samples were then centrifuged at 3700 RPM, and the water:MeOH phase was transferred to 2 mL Eppendorf tubes and stored at −20 °C until further treatment (see below).

For the concurrent extraction of metabolites and lipids, the procedure was modified to use a water:methanol:DCM (1:1:2) mixture as the solvent. The samples were sonicated in a Bandelin Sonorex water bath and rested on ice to allow phase separation. The MeOH:water phase was treated as before, and the DCM phase was transferred to Eppendorf tubes, stored at −20 °C until derivatization and analysis.

### 5.3 Derivatization of metabolites and gas chromatography

Metabolites were analyzed using a modified version of the derivatization protocol described by Sogin et al. (2019). The cultures used in this study are listed in Table 1. Samples were further dried by adding 250 µL of toluene, sonication, and drying using a rotary evaporator (Concentrator Plus, Eppendorf). Due to the high salt concentrations in the supernatant samples, they were dried for an additional 12 hours. Toluene was evaporated at 37°C at 250 RPM, and 80 µL of methoxyamine dissolved in dry pyridine (20 mg mL^−1^) was added to the dry sample, which was ultrasonicated for 10 minutes. Samples were then incubated at 37°C for 90 minutes at 1350 RPM to evaporate the pyridine. Then, 100 µL N,O-bis(trimethylsilyl)trifluoroacetamide (BSTFA) was added. This mixture was sonicated and incubated at 37°C for 30 minutes, then centrifuged at 13,000 rpm for 2 minutes. The clear supernatant was analyzed using an Agilent 7890B GC coupled to an Agilent 5977A single-quadrupole mass-selective detector. 1 μl sample was injected via the GC inlet liner (ultra inert, splitless, single taper, glass wool; Agilent) onto a DB-5MS column (30 m by 0.25 mm, 0.25-μm film thickness, including 10 m DuraGuard column; Agilent). The inlet liner was changed every 100 samples to prevent damage to the GC column and associated retention-time shifts. The injector temperature was set at 290°C. Chromatography was performed with an initial column oven temperature set at 60°C, followed by a ramp of 20°C min^−1^ to 325°C, then held for 2 min. Helium carrier gas was used at a constant flow rate of 1 ml min^−1^. Mass spectra were acquired in electron ionization mode at 70 eV across the mass range m/z 50-600 at a scan rate of 2 s^−1^. The retention time for the method was locked using a standard mixture of fatty acid methyl esters (Sigma-Aldrich). Peaks were annotated using Agilent Unknowns with peak filters of 28, 91, and 149, with area filters set to greater than or equal to absolute counts of 10. Deconvolution settings were an RT window size factor of 400, an extraction window of 0.3 m/z to the left and 0.7 m/z to the right, a sharpness threshold of 25%, a minimum number of ion peaks of 3, and a maximum number of peak shapes to store of 10. Spectral search for compound identification was done against the NIST/EPA/NIH Mass Spectral Database (NIST 11), where match factors, scaled from 0 to 99, reflect the similarity between the observed spectrum and reference entries. Values above 90 are generally considered high-confidence matches.

To quantify the total amount of soluble protein, extraction was performed as before, except using 5 mL of MeOH:water. Of these, 2 mL were used for metabolite extraction as before, and 2 mL were used for protein quantification with the Thermo Fisher MicroBCA kit according to the manufacturer’s instructions. The ratio R of metabolite *i* in sample *k* (µg metabolite per µg of protein) was then calculated as follows:

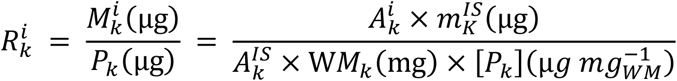

Where 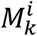 is the metabolite mass of metabolite *i* and *P*_*k*_ is the soluble protein mass in sample *k*, respectively. 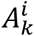 and 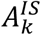 are the peak areas of metabolite *i* and the ribitol internal standard in sample *k*, respectively. *WM*_*k*_(mg) is the total wet mass of sample *k* from which both the metabolite extraction and protein extractions were derived, and [*P*_*k*_] is the concentration of protein per mg of wet mass of sample *k*. Each GC run was accompanied by a calibration mix composed of 2.5 µg of Lauric acid, 2-hydroxyhexanoic acid, Sucrose, Trehalose dihydrate, DL-Lysine, Leucine, Glycine, Di-sodium succinate hexahydrate, L-Ornithine monohydrochloride, DL-Serine, L-Asparagine, DL-Tryptophane, and Cholesterol.

### 5.4 Curation, annotation, and phylogenetic analyses of Halobacteriota and Desulfobacterota genomes from the GTDB

The GTDB accessions of the phyla Halobacteriota and Desulfobacteriota were fetched with the gtt-get-accessions-from-GTDB command from the GTotree (v1.8.3) pipeline (Chklovski et al., 2023; Lee, 2019). From this group of datasets, all known entries of alkanotrophic archaea except ANME-3 were subsetted based on the literature, with ANME-1 and ANME-2 defined as in Chadwick et al., 2022, and Laso-Pérez et al., 2023, *Ca*. Ethanoperedens as defined by Chen et al. 2019 and Hahn et al. 2020, *Ca*. Syntropharchaeum, as defined by Laso-Pérez et al. 2016, *Ca*. Alkanophaga, as defined by Zehnle et al. 2023 and *Ca*. Methanoliparia is defined as by Laso-Pérez et al., 2019. In addition, a custom subset of non-ANKA Halobacteriota was randomly selected, with the criterion that no individual genus could be represented by more than 5 entries to prevent overrepresentation of lab strains of the Halobacteriales.

For Desulfobacteria, similarly, all GTDB entries that match the Seep-SRB-1a, -1d, and -1g clades, the Seep-SRB2 clade, and the class Desulfofervidia were selected, according to Schreiber et al. 2010 and Murali et al. 2023. In addition, a curated subset of Desulfovibrionia, Desulfuromonadia, and Desulfobacterales, excluding the known ANKA partner bacteria, was randomly selected using the same genus criteria as above, along with the additional criterion that no single order accounts for more than 20% of the total dataset. This prevented the overrepresentation of the very abundant Desulfovibrionales genomes. The accession to all genomes analyzed in this study and their GTDB metadata are provided in Supplementary Table 6.

### 5.5 Annotation of genes for disaccharide metabolism and protein phylogenies

A set of curated HMMs of various protein families associated with trehalose, sucrose, and general disaccharide metabolism was downloaded from Interpro (v.104.0) (Blum et al., 2025). The profiles used, their parent families, and associated functions are listed in Table 2. These were searched against the curated sets of GTDB genomes using *hmmsearch* (v.3.4) (Eddy, 2011). Hmmsearch outputs were parsed using a custom Python script, and hits were filtered for E-values ≤ 1e-10 and hmm coverage ≥ 60%. To prevent cross-annotation, each locus tag was only considered for its best hit.

The amino acid sequences of TreT, TreP/MalP, and OtsA were analyzed for phylogeny. We extracted the sequences of locus tags with valid HMM hits from the Prokka annotations of the genomes(Seemann, 2014). For each HMM profile, only the locus tag associated with the best hit was considered. These were then concatenated and aligned using MAFFT-Linsi (v7.520) (Katoh et al., 2005; Katoh and Standley, 2013) and trimmed using trimAL (v1.4) (Capella-Gutiérrez et al., 2009) with the *-gappyout* option. The trimmed alignments were passed to IQtree2 with the automatic model finder and 1000 ultrafast bootstraps(Kalyaanamoorthy et al., 2017; Minh et al., 2020). All trees were visualized and edited with ITOL(v7)(Letunic and Bork, 2024).

### 5.6 Assimilation of ^13^C-DIC and ^13^C-trehalose into bacterial and archaeal lipids

We assessed the dependence of CO_2_ assimilation and trehalose assimilation on alkane availability using the readily available Ethane50 culture (Hahn et al., 2020). Therefore, we washed the Ethane50 biomass from two 700-ml bottles and transferred 25 mL of dense Ethane50 biomass into replicate 150-ml culture bottles, which we filled to 100 mL with 10 mM DIC-containing culture medium.

For the ^13^C-DIC labeling experiment, 6 subcultures of Ethane50 received 100 µl of a ^13^C-DIC standard solution (1 NaH^13^CO_3_; Sigma, in MilliQ water), resulting in δ^13^C-DIC values of +550‰. These subcultures were split into two sets of 3 biological replicates; half of the replicates were treated with ethane, and the other half received no exogenous energy source. Sulfide was sampled after 0,3,6,9, and 24 days of incubation, and the first and last time points were sampled for lipids and DIC.

For the ^13^C-trehalose experiment, an anoxic stock of 20 mg mL^−1^ 5% ^13^C-labelled trehalose was prepared by adding 10 mg of ^13^C-labeled trehalose (Sigma-Aldrich) and 190 mg of non-labeled trehalose (Merck) to 10 mL of anoxic water. The stock was sterile-filtered (0.2 µm, Minisart, Sartorius) and stored under oxygen-free conditions. Due to the limited availability of biomass from these sediment-free enrichment cultures derived from deep-sea sediments, we prioritized time-series resolution over replicate numbers. As described above for the DIC incubation, 11 subcultures of Ethane50 were prepared; however, they were not labeled with DIC. Instead, 10 of the subcultures received 100 µL of the ^13^C-trehalose stock solution. One subculture was kept trehalose-free as a control. The 10 trehalose-containing subcultures were then split into two experimental sets of five, with one set given ethane as an alkane energy source and the other kept on an N2:CO2 gas mixture. The cultures were incubated at 50 °C for 15 days; at 0, 3, 6, 9, and 15 days, sulfide was measured from all remaining bottles, and one culture bottle was sacrificed for lipid extraction in each experimental condition.

For fatty acid analysis, one-third of the total lipid extract (TLE) was saponified with 6% potassium hydroxide (Sigma-Aldrich) in methanol (MeOH, Sigma-Aldrich) for 3 h at 80°C. The product was extracted three times with n-hexane under basic conditions to obtain neutral lipids (NL), followed by hexane extraction after acidification with HCl, yielding the fatty acids (FA). The FA fraction was dried and derivatized with 10% BF3 in MeOH (VWR) at 70°C for 1 h to form fatty acid methyl esters (FAMEs), which were then extracted with hexane. To yield the isoprenoids from the archaeal ether lipids, one-third of the TLE was subjected to ether cleavage using the boron tribromide treatment, followed by reduction with lithium triethylborohydride (Summons et al., 1998).

Analyses of FAMEs and ether-cleaved isoprenoids were carried out using gas chromatography (GC) coupled to a flame ionization detector (Trace GC Ultra, Thermo Scientific). GC cross-checked compound identities–mass spectrometry (Trace 1310 + ISQ 700, Thermo Scientific) operated in EI mode at 70 eV and collecting full-scan spectra from m/z 50 to 850. Stable carbon isotope compositions (expressed as δ^13^C values relative to VPDB) were determined using GC–isotope ratio mass spectrometry (Trace GC Ultra connected with a GC-IsoLink and interfaced through a ConFlo IV to a Delta V Plus IRMS, Thermo Scientific). All GC analyses were performed in splitless mode injection (1 min at 310°C) and used an identical Restek Rxi-5 ms column (30 m × 0.25 mm ID × 0.25 μm, Restek). The temperature program for all instruments was: 60°C for 1 min, followed by heating to 150°C at 10°C min^−1^, then to 310°C at 4°C min^−1^, with a final hold at 310°C for 30 min for the previously cleaved isoprenoids from the archaeal lipids, and 20 min for FAMEs. Helium served as the carrier gas at 1.0 mL min^−1^, except during GC–IRMS runs where 1.2 mL min^−1^ was used. For the latter analyses, compounds eluting from the column were oxidized to CO_2_ in a combustion reactor maintained at 940°C. The instrument was calibrated against a CO_2_ reference gas with a known isotopic composition, and precision was monitored by repeated analysis of a C20–C40 n-alkane mixture, which exhibits long-term variability of ±0.5‰. δ^13^C values of FAMEs were corrected for the added carbon introduced during derivatization.

## 6 Author contributions

ST and GW conceived of the study, with input from all authors. ST extracted, derivatized, and analyzed samples with GC-MS with support from ML. LS derivatized and analyzed lipids under the supervision of ME. ST performed bioinformatic work. ST and GW wrote the manuscript. All authors provided critical insights.

## 7 Acknowledgements

The authors thank Anja Seebeck and Susanne Menger for maintaining the alkane-oxidizing cultures. We thank Adrian Stolfa and Kristina Weinert for critical support in sample preparation and mass spectrometry. Dr. Grace D’Angelo provided valuable insights into sugar metabolism and metabolomics. This research was enabled by funding from the Hector Fellow Academy for the PhD project of Stian Torset, and by the Deutsche Forschungsgemeinschaft (DFG) through Germany’s Excellence Initiative Cluster of Excellence EXC 2077 ‘The Ocean Floor—Earth’s Uncharted Interface’ (EXC-2077-390741603)

