## Supplementary Information for "Alkane-oxidizing consortia produce substantial amounts of disaccharides"

Supplementary Material

### 1 Supplementary Figures and Tables

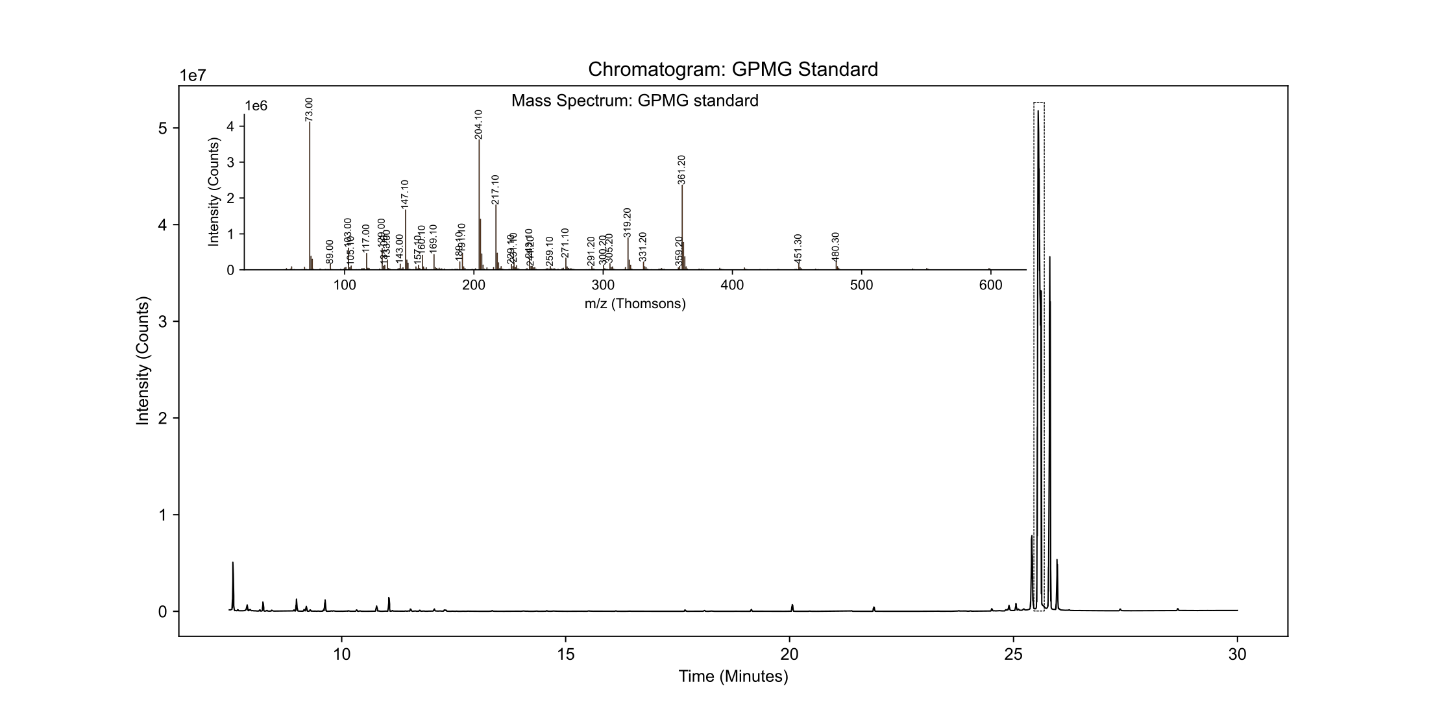

Figure S1: Chromatogram and Mass spectra of the Galactopyranosyl-D-mannopyranose (GPMG) standard. Inset on the chromatogram is the Mass spectra corresponding to the peak highlighted by a dashed box.

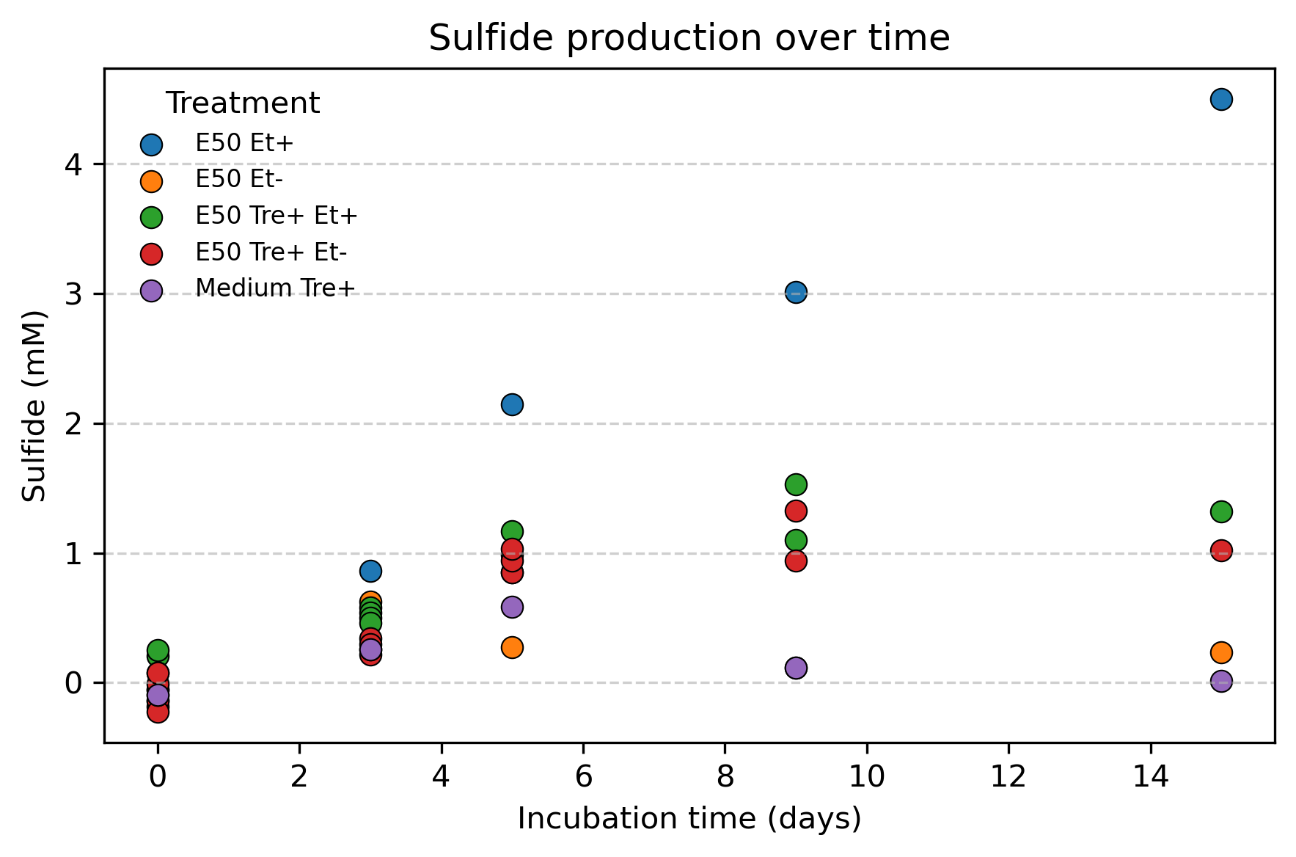

Fig S2: Sulfide production measured over time in incubations of Ethane50 with or without ethane and with and without trehalose, including a medium blank with trehalose.

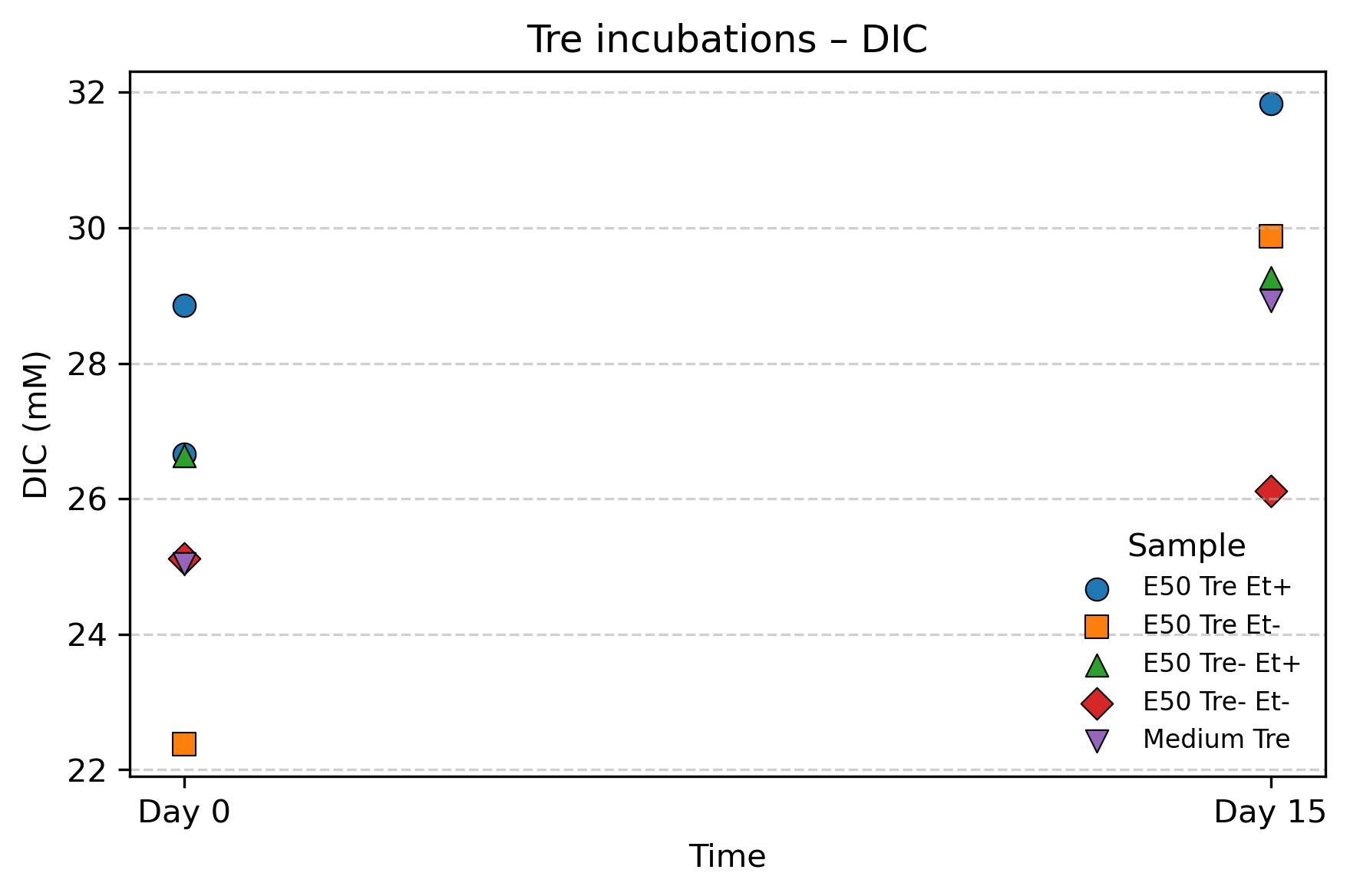

FigS3: Medium DIC concentrations at the start and end of incubations for Ethane50 enrichment cultures incubated with or without ^13^C labelled trehalose.

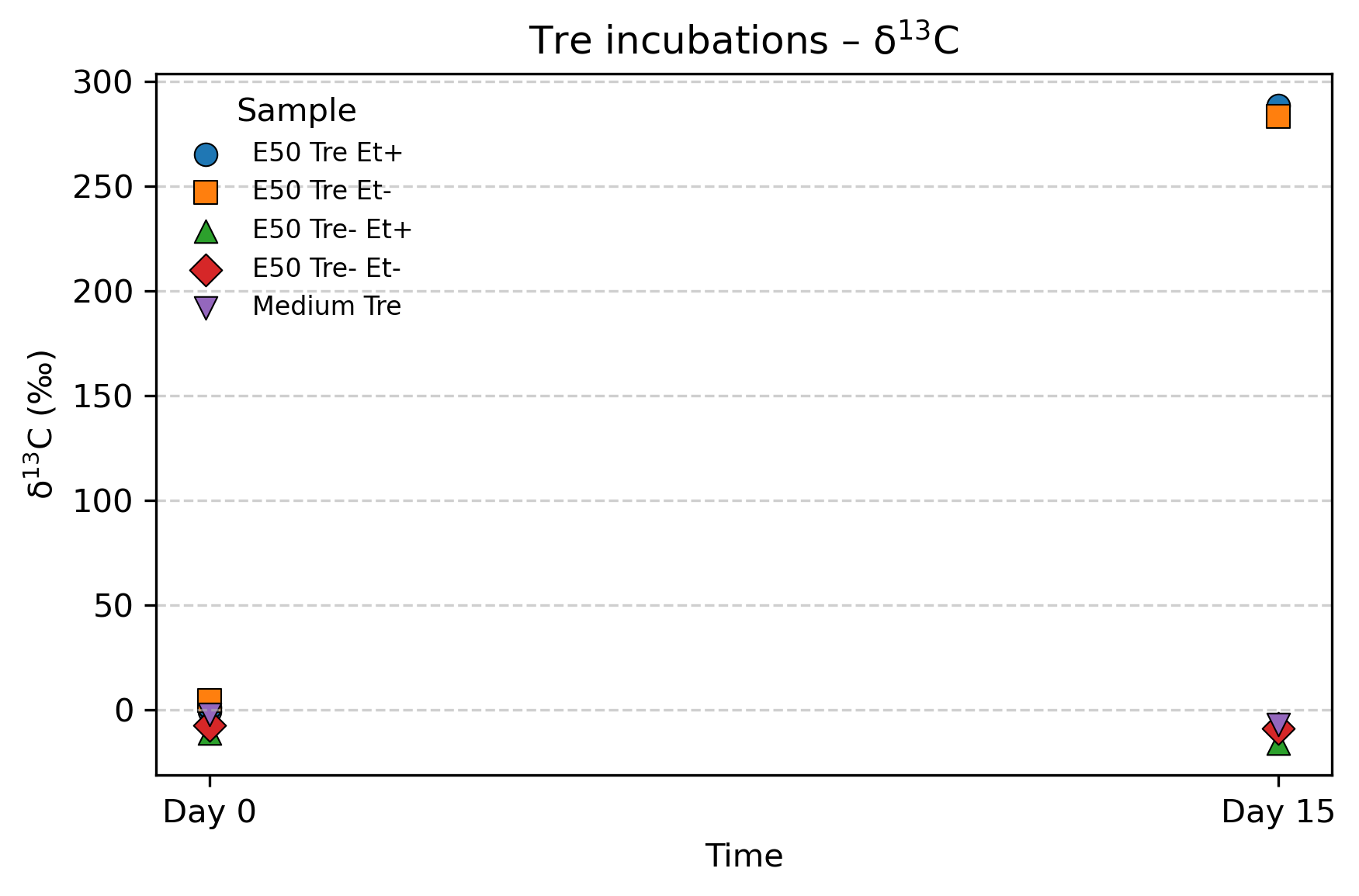

FigS4: δ^13^C values for the medium DIC at the start and end of incubations for Ethane50 enrichment cultures incubated with or without ^13^C labelled trehalose.

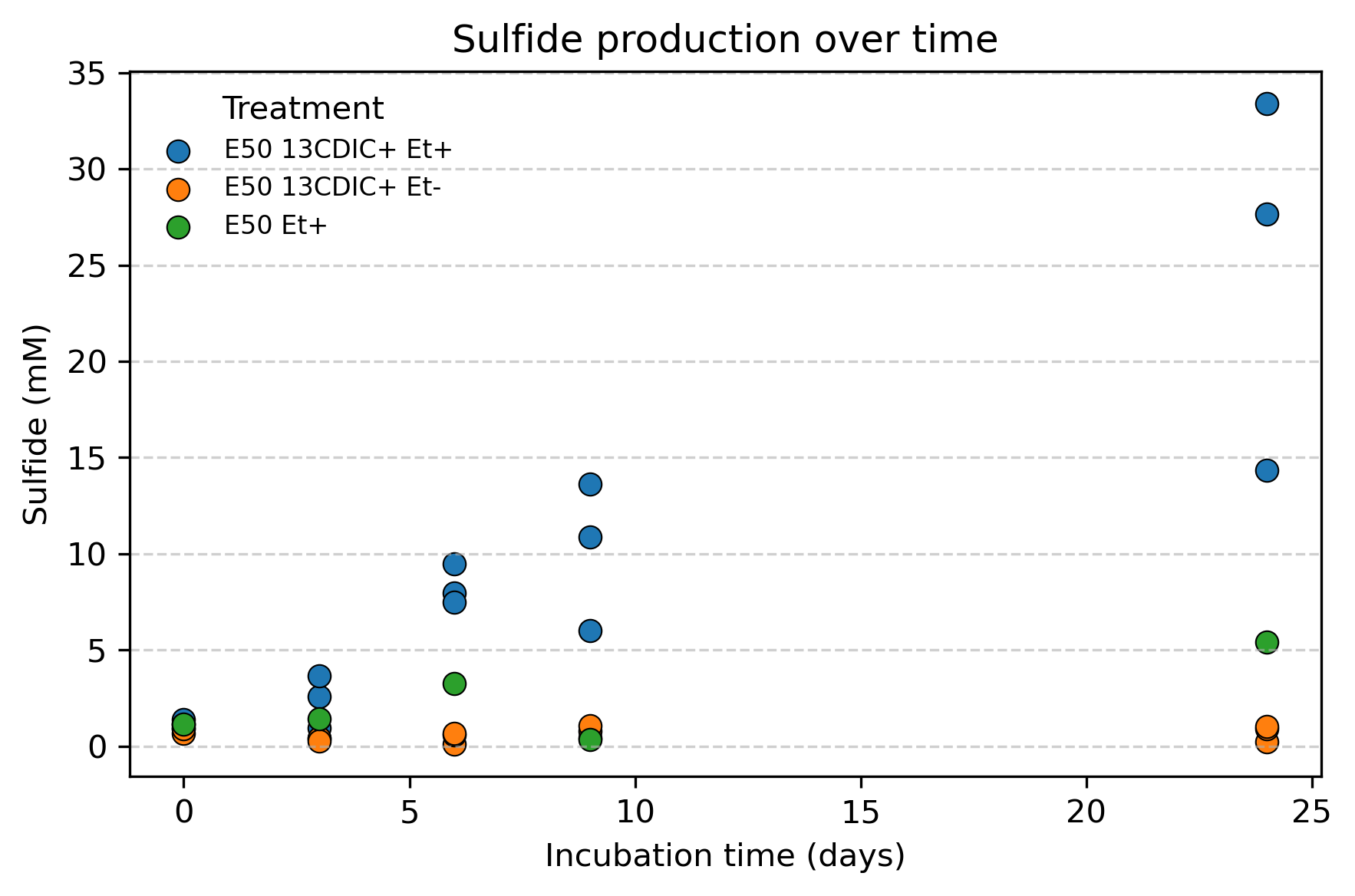

Figure S5: Sulfide production in Ethane50 enrichment cultures incubated with labelled DIC with or without ethane.

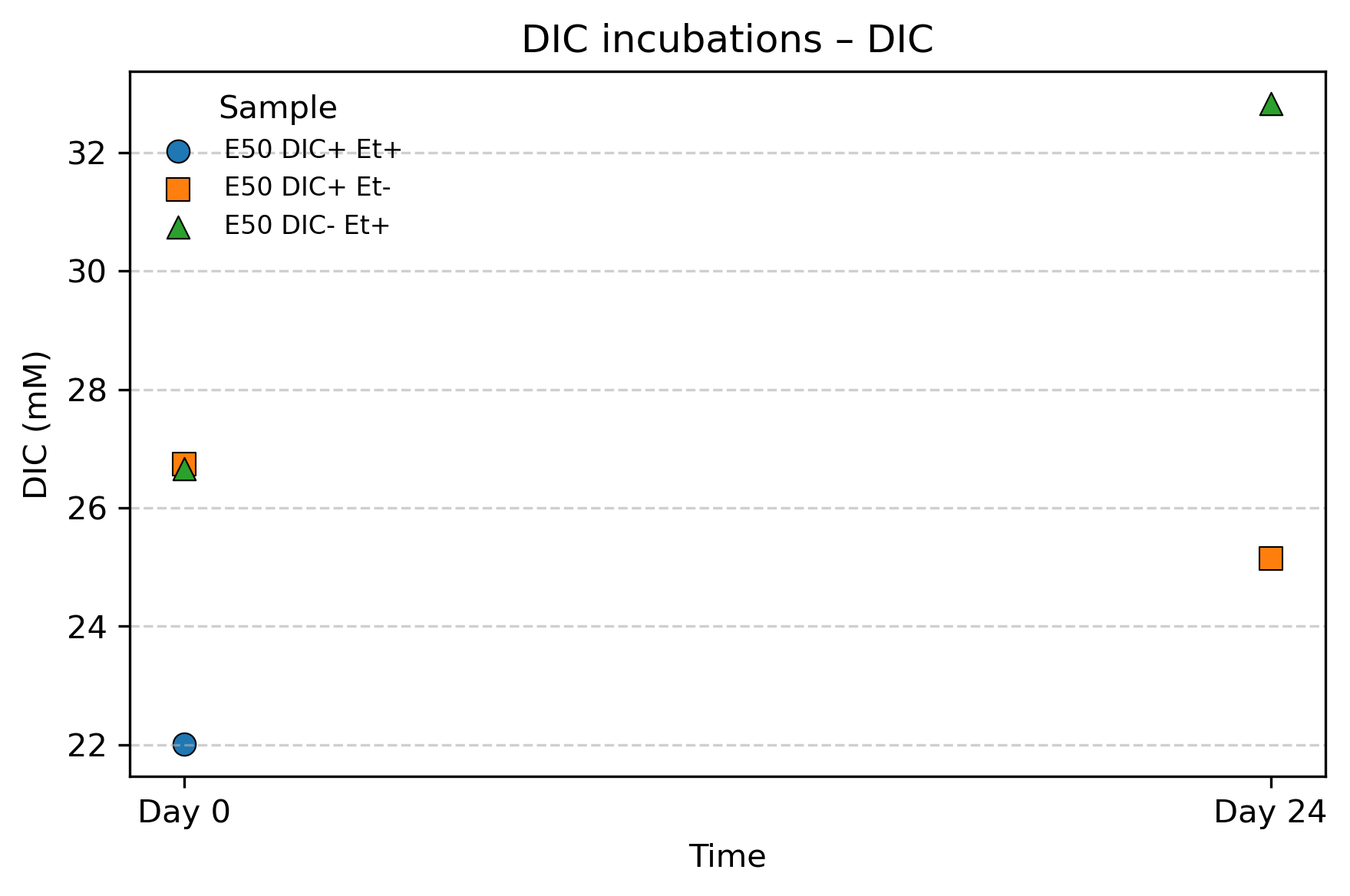

Figure S6: DIC concentrations in medium of Ethane50 cultures incubated with or without labelled DIC and with or without ethane. Note that the DIC+ Et+ sample was lost due to an equipment failure.

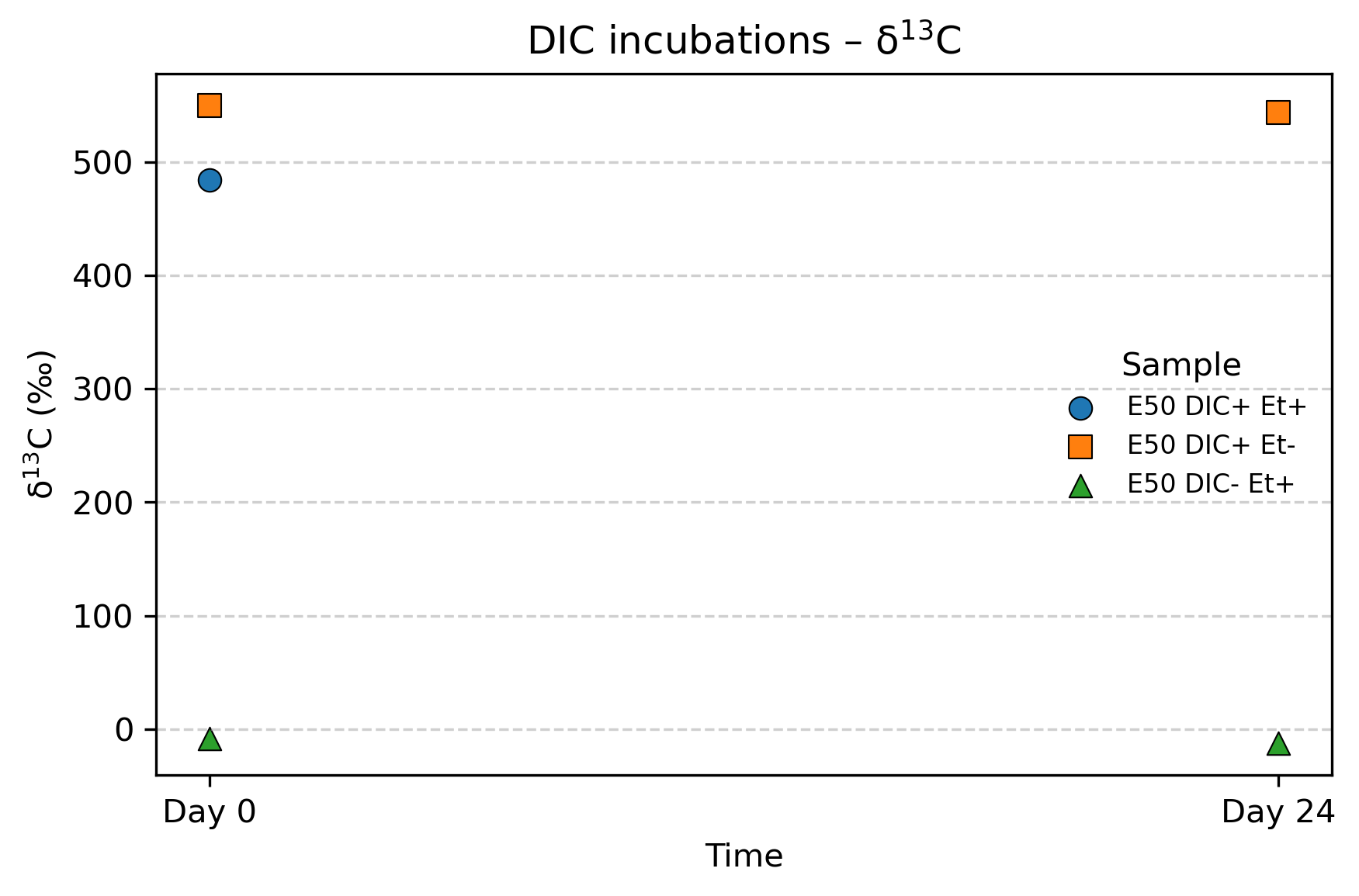

FigS7: δ13C values for the medium DIC at the start and end of incubations for Ethane50 enrichment cultures incubated with or without 13C labelled DIC.
